# Phagosome-mediated activation of STING during diverse bacterial infections

**DOI:** 10.1101/2022.10.28.514268

**Authors:** Stephanie A. Ragland, Jonathan C. Kagan

## Abstract

Central to health and disease are innate immune receptors that bind bacterial molecules to initiate inflammation and host defense. Beyond pathogens and their membrane disruptive activities, mechanisms enabling bacterial molecules to access innate immune receptors in the cytoplasm are unknown. Here, we describe the cytoplasmic cyclic dinucleotide (CDN)-binding protein STING as a common bacterial sensor. Irrespective of virulence and after bacteriolysis in phagolysosomes, CDNs produced during infections with evolutionarily diverse bacteria activate STING. Of the several known CDN transporters, two supported bacteria-induced STING activation. We propose a connection between phagocytosis and STING that ensures host-bacteria interactions result in STING activation.

---

Innate immune pattern recognition receptors (PRRs) induce inflammation upon binding microbial molecules known as pathogen-associated molecular patterns (PAMPs). Several PRR families exist, such as the Toll-like receptors (TLRs) whose PAMP-recognition domains survey the extracellular and intra-endosomal space (*1*). Other PRRs survey the cytosol, including the RIG-I-like receptors (RLRs), the nucleotide-binding domain leucine-rich repeat containing proteins (NLRs), cyclic GMP-AMP synthase (cGAS), and stimulator of interferon genes (STING) (*2-4*). While PAMP detection is a requisite aspect of PRR-mediated host defense, our understanding of how PRRs access microbial ligands is incomplete. This lack of knowledge is most notable in the context of cytoplasmic PRRs, whose access to microbes and their products is restricted by the plasma and endomembranes of the cell. The means by which PAMPs cross membranes to access cytoplasmic PRRs have largely been studied in the context of pathogenic bacteria that induce phagosomal membrane damage. As such, cytoplasmic PRRs are often framed as pathogen-specific receptors (*5, 6*). TLRs, in contrast, due to their relative ease of PAMP access in the extracellular and luminal endosomal environments, are considered the only family of PRRs that function as general sensors of bacterial encounters.

Despite this prevailing view, a few studies reported that cytoplasmic PRRs can detect non-pathogens, including *Escherichia coli, Mycobacterium smegmatis*, and *Listeria innocua (7-9)*. It is unknown whether cytoplasmic PRRs are general contributors to host-bacteria interactions, nor do we know how cytoplasmic PRRs access PAMPs produced by most bacteria. A growing body of literature suggests that protein transporters facilitate translocation of specific molecules (*e*.*g*., nutrients, peptides, and PAMPs) from the endosomal lumen to the cytosol (*10-12*). For example, PAMPs that stimulate the NLRs NOD1 and NOD2 are endocytosed and then translocated into the cytosol by one of several transporters (*13-15*). Cyclic dinucleotides (CDNs) that stimulate STING are also translocated into the cytoplasm by several transporters (*16-20*). However, as these transporters have been studied primarily in the context of chemically-synthesized and/or highly purified PAMP molecules, we have limited knowledge for their role in cellular encounters with actual microorganisms.

In this study, we report that, of all the cytoplasmic PRRs, STING is activated by the widest spectrum of evolutionarily diverse bacteria. Bacteria-induced STING activation occurs regardless of virulence potential but requires phagocytosis and intra-phagosomal bacterioloysis. Two previously identified CDN transporters, SLC46A2 and LRRC8A (*18-20*), supported STING activation by diverse intra-phagosomal bacteria producing CDNs. These findings explain how phagocytosis is coordinated with innate immune stimulation and establish STING as a common sensor of bacteria-host interactions.

## Results

### Evolutionarily diverse bacteria stimulate TLR-independent interferon (IFN) responses in macrophages

To identify common innate immune activities in macrophages, we assembled a panel of evolutionarily diverse bacteria consisting of gram-positive bacteria (*Staphylococcus aureus, Staphylococcus epidermidis, L. innocua, Lactococcus lactis, Lactobacillus plantarum, Micrococcus luteus, Enterococcus faecalis*, and *Bacillus subtilis*) and gram-negative bacteria (*E. coli, Neisseria perflava, Yersinia pseudotuberculosis, Bacteroides fragilis*, and *Bacteroides thetaiotaomicron*) (**Fig. 1A**). We performed synchronous, side-by-side infections of wild-type (WT) primary murine bone marrow-derived macrophages (BMDMs) and measured the production of the inflammatory mediators TNFα, interferon β (IFNβ), and IP-10. By 24 hours post-exposure, all bacteria induced the secretion of TNFα, IFNβ, and IP-10 (**Fig. 1B**). For IFNβ and IP-10, which are constituents of the IFN response pathway, their production was stimulated by gram-negative bacteria within 4 hours of infection whereas most gram-positive bacteria induced these responses within 24 hours (**Fig. 1B**). *Bacteroides* species stimulated weak IFN responses and TNFα secretion, compared with other gram-negatives (**Fig. 1B**).

**Figure 1.**
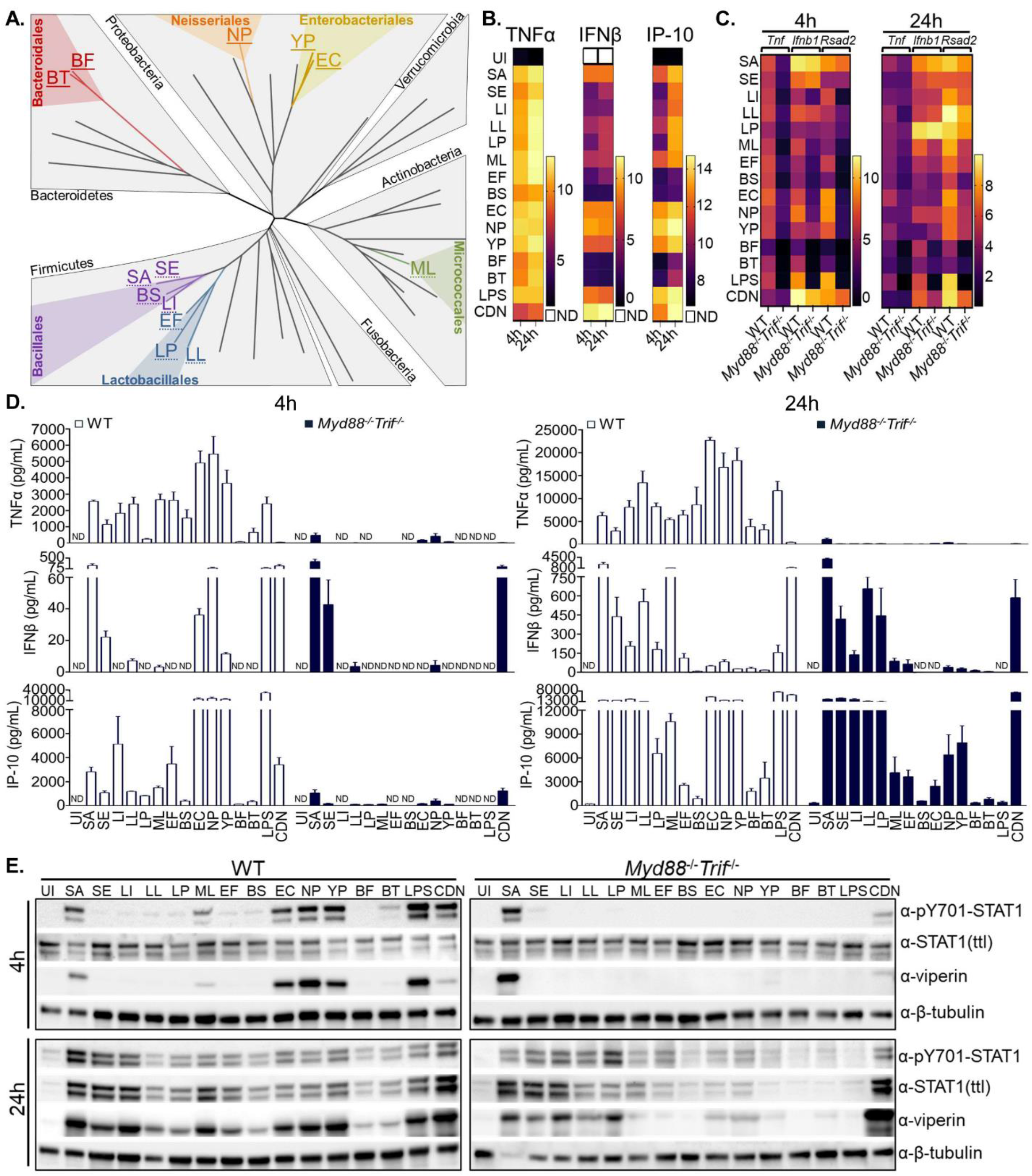
Evolutionary diverse bacteria stimulate TLR-independent IFN responses in macrophages. **(A)** Phylogenetic tree of the most common bacterial phyla associated with humans, compiled from species representing 28 orders. Bacteria used in this study are distinguished by colored text, and bacteria of the same colored text are within the same taxonomic order. Gram-negative bacteria are denoted with a solid underline whereas gram-positive bacteria are denoted with a dashed underline. Abbreviations: *S. aureus* (“SA”), *S. epidermidis* (“SE”), *L. innocua* (“LI”), *L. lactis* (“’LL”), *L. plantarum* (“LP”), *M. luteus* (“ML”), *E. faecalis* (“EF”), *B. subtilis* (“BS”), *E. coll* (“EC”), *N. perflava* (“NP”), *B. fragilis* (“BF”), *B. thetaiotaomicron* (“BT”), and *Y. pseudotuberculosis* (“YP”). **(B)** WT BMDMs were untreated (or uninfected, herein “Ul”) or exposed to 1 pg/mL LPS or ADU-S100 (“CDN”) and bacteria at an MOI of 50. At 4 h and 24 h post-exposure, conditioned supernatants were analyzed by ELISA for TNFα (left), IFNβ (middle), and IP-10 (right) protein abundance. Heat map values represent Log_10_ protein abundance. White boxes indicate protein abundance below the limit of detection (“not detected", herein abbreviated “ND”). **(C)** WT and *Myd88*^-/-^TriF^-/-^ iBMDMs were treated as in **Fig. 1B**. 4 h and 24 h RNA lysates were analyzed for *Tnf, Ifnbl*, and *Rsad2* transcript abundance by qRT-PCR with respect to the tbp internal control, and fold change was calculated relative to Ul control. Heat map values are fold change expressed as Log_10_. **(D)** 4 h (left) and 24 h (right) conditioned supernatants from **Fig. 1C** were analyzed by ELISA for TNFα (top), IFNβ (middle), and IP-10 (bottom) protein abundance. **(E)** 4 h (top) and 24 h (bottom) whole cell protein lysates from **Fig. 1C** were analyzed by immunoblot for phosphorylated STAT1 (pY701), total STAT1, viperin, and β-tubulin.

To corroborate these findings, similar studies were performed in WT immortalized BMDMs (iBMDMs). In response to bacteria, we observed transcriptional responses for *Ifnb1* and the IFN stimulated gene (ISG) *Rsad2*, secretion of IFNβ and IP-10, the phosphorylation of the IFN-responsive transcription factor STAT1 (herein “pSTAT1”), and the production of the *Rsad2* product viperin from iBMDMs (**Fig. 1C-E**). The relative kinetics and magnitude of all IFN responses examined was similar to those observed in primary BMDMs (**Fig. 1B**). Similar trends were made with *Tnf* transcriptional responses and TNFα protein abundance in iBMDMs (**Fig. 1C, D**).

To determine the role of TLRs in macrophage responses to these diverse bacteria, we employed iBMDMs doubly deficient in the adaptor proteins MyD88 and TRIF. These cells are defective for all TLR-dependent inflammatory activities (*21*). As expected, pure TLR ligands failed to stimulate the transcription of *Tnf* or the production of pSTAT1 and viperin in *Myd88*^*-/-*^*Trif*^*-/-*^ iBMDMs (**Supplemental Fig. 1A, B**). In contrast, *Tnf*, pSTAT1, and viperin responses to ligands for the cytoplasmic PRRs STING and NOD2 were intact in *Myd88*^*-/-*^*Trif*^*-/-*^ iBMDMs (**Supplemental Fig. 1A, B**). Compared to WT cells, *Myd88*^*-/-*^*Trif*^*-/-*^ iBMDMs were defective for TNFα secretion in response to all bacteria examined (**Fig. 1D**), indicating that TLRs are the primary drivers of TNFα production during diverse bacterial encounters.

All bacteria we examined induced IFN responses by 24 hours in *Myd88*^*-/-*^*Trif*^*-/-*^ cells (**Fig. 1C-E**). A multiplicity of infection (MOI) as low as 1 was sufficient for most bacteria to stimulate an increase of pSTAT1, total STAT1, and viperin protein abundance in WT and *Myd88*^*-/-*^*Trif*^*-/-*^ iBMDMs, with *Bacteroides* species being the weakest inducers (**Supplemental Fig. 1C**). IFN responses were not associated with host cell death, as lactate dehydrogenase release, a marker for inflammatory cell death, did not correlate with IFN activities induced by the collection of bacteria examined (**Supplemental Fig. 1D**). These findings demonstrate that evolutionarily diverse bacteria stimulate TLR-independent IFN responses.

### TLR-independent IFN responses to bacteria require a TBK1-IRF3-IFNAR signaling axis

We defined the pathway features important for TLR-independent IFN signaling using a refined group of bacteria. Since we observed STAT1 activation in *Myd88*^*-/-*^*Trif*^*-/-*^ iBMDMs exposed to bacteria (**Fig 1E, Supplemental Fig. 1C**), we hypothesized a role for the IFN-α/β receptor (IFNAR) in TLR-independent IFN signaling. We used CRISPR/Cas9 to inactivate *Ifnar1* in *Myd88*^*-/-*^*Trif*^*-/-*^ iBMDMs. As expected, recombinant IFNβ protein was unable to elicit IP-10 secretion from *Myd88*^*-/-*^*Trif*^*-/-*^*Ifnar1*^*-/-*^ iBMDMs (**Supplemental Fig. 1E**). Upon exposure to the examined bacteria, IP-10 protein abundance was also reduced in *Myd88*^*-/-*^*Trif*^*-/-*^ *Ifnar1*^*-/-*^ iBMDMs, compared with *Myd88*^*-/-*^*Trif*^*-/-*^ cells (**Supplemental Fig. 1E**). All genotypes of iBMDMs exhibited a similar capacity to secrete IFNβ in response to bacteria (**Supplemental Fig. 1F**). Thus, IFNAR is required for TLR-independent ISG production, but not IFNβ production, in response to bacterial encounters.

Most cytoplasmic PRRs signal through IRF3 to promote IFN transcription (*6*). We found that exposure of *Myd88*^*-/-*^*Trif*^*-/-*^ iBMDMs to bacteria resulted in increased abundance of activated IRF3 (*i*.*e*., IRF3 phosphorylated at S396) (**Supplemental Fig. 1G**). To determine if IRF3 is required for IFN responses to bacteria, we generated *Irf3*-deficient *Myd88*^*-/-*^*Trif*^*-/-*^ iBMDMs (**Supplemental Fig. 1H**). Upon exposure to a modified CDN, named “ADU-S100” (a synthetic bacterial CDN analog, 2′3′-c-di-AM(PS)2 (Rp, Rp)) that binds STING to activate IRF3, *Myd88*^*-/-*^*Trif*^*-/-*^*Irf3*^*-/-*^ iBMDMs were defective for IP-10 secretion, compared to *Myd88*^*-/-*^*Trif*^*-/-*^ cells (**Supplemental Fig. 1I**). In contrast, IRF3 was dispensable for IP-10 production in response to recombinant IFNβ protein (**Supplemental Fig. 1I**). We found that all bacteria examined were defective for inducing IP-10 secretion from *Myd88*^*-/-*^*Trif*^*-/-*^*Irf3*^*-/-*^ iBMDMs, indicating that IRF3 is required for TLR-independent IFN production to bacteria (**Supplemental Fig. 1I**).

IRF3 is commonly activated by the kinase TBK1 (*6*). We observed increased abundance of activated TBK1 (*i*.*e*., TBK1 autophosphorylated at S172, herein “pTBK1”) in *Myd88*^*-/-*^*Trif*^*-/-*^ iBMDMs exposed to bacteria (**Supplemental Fig. 1J**). We generated *Tbk1*-deficient *Myd88*^*-/-*^*Trif*^*-/-*^ iBMDMs (**Supplemental Fig. 1K**) and found that *Myd88*^*-/-*^*Trif*^*-/-*^ *Tbk1*^*-/-*^ iBMDMs were defective for IP-10 secretion upon exposure to ADU-S100 or bacteria, compared with *Myd88*^*-/-*^*Trif*^*-/-*^ iBMDMs (**Supplemental Fig. 1L**). Secretion of IP-10 in response to recombinant IFNβ was similar between *Tbk1*-deficient or -proficient cells (**Supplemental Fig. 1L**). Together, these data indicate that a common feature of the TLR-independent IFN response to bacteria is the dependence on a TBK1-IRF3-IFNAR signaling axis.

### STING controls TLR-independent IFN responses to bacteria in macrophages

To determine which cytoplasmic PRR mediates bacteria-induced IFN responses, we examined proteins involved in activating TBK1-IRF3-IFNAR signaling. MAVS is an adaptor protein required to induce IFN responses downstream of the RLRs (*2*). We generated *Myd88*^*-/-*^*Trif*^*-/-*^*Mavs*^*-/-*^ iBMDMs (**Supplemental Fig. 1M**) and observed that *Mavs*-deficiency ablated IP-10 production upon exposure to RLR agonistic Sendai virus (**Supplemental Fig. 1N**). In contrast, MAVS was not required for IP-10 secretion upon exposure to ADU-S100 and all bacteria examined (**Supplemental Fig. 1N**). MAVS is therefore not responsible for TLR-independent IFN responses to bacteria.

The peptidoglycan sensor NOD2 reportedly signals through the RIP2 kinase to stimulate IFN responses via TBK1 and IRF3 (*22*). We therefore generated *Myd88*^*-/-*^*Trif*^*-/-*^*Rip2*^*-/-*^ iBMDMs (**Supplemental Fig. 1O**). We found that compared to parental cells, *Rip2*-deficiency resulted in ablated IP-10 responses to the NOD2 agonist N-glycolyl-muramyl dipeptide but normal IP-10 responses to ADU-S100 (**Supplemental Fig. 1P**). *Myd88*^*-/-*^*Trif*^*-/-*^*Rip2*^*-/-*^ iBMDMs exhibited modest reductions in IP-10 in response to bacterial treatments, compared to *Myd88*^*-/-*^*Trif*^*-/-*^ iBMDMs (**Supplemental Fig. 1P**). To further investigate the role of RIP2 during infection, we examined early *S. aureus*-induced IFN responses and found that pTBK1, pSTAT1, and viperin protein abundance was comparable between *Myd88*^*-/-*^*Trif*^*-/-*^ and *Myd88*^*-/-*^*Trif*^*-/-*^*Rip2*^*-/-*^ iBMDMs (**Supplemental Fig. 1Q**). This finding suggests that RIP2 may contribute to sustained IFN responses but is not required for initiation of these responses.

We next examined a role for STING. STING detects CDNs produced by bacteria but can also detect 2′3′-cGAMP (herein “2′3′-cGA”), which is synthesized by cGAS upon cGAS recognition of cytosolic DNA (*3, 23*). We found that *Myd88*^*-/-*^*Trif*^*-/-*^ iBMDMs exposed to bacteria exhibited an increase in the abundance of activated STING (*i*.*e*., STING phosphorylated at S365, herein “pSTING”) (**Fig. 2A**). We therefore generated *Myd88*^*-/-*^*Trif*^*-/-*^*Sting*^*-/-*^ iBMDMs (**Supplemental Fig. 2A**). *Myd88*^*-/-*^*Trif*^*-/-*^ and *Myd88*^*-/-*^ *Trif*^*-/-*^*Sting*^*-/-*^ iBMDMs secreted similar amounts of IFNβ and IP-10 in response to Sendai Virus infection (**Fig. 2B, Supplemental Fig. 2B**). In contrast, *Myd88*^*-/-*^*Trif*^*-/-*^ *Sting*^*-/-*^ iBMDMs were defective for IFNβ and IP-10 secretion in response to agonists that stimulate STING directly (the modified CDN, ADU-S100) or indirectly (cytosolic DNA via cGAS) (**Fig. 2B, Supplemental Fig. 2B**). Exposure of cells to every species of bacteria in our panel revealed an ablation of IFNβ and IP-10 secretion in *Myd88*^*-/-*^*Trif*^*-/-*^*Sting*^*-/-*^ iBMDMs, compared to *Myd88*^*-/-*^*Trif*^*-/-*^ cells (**Fig. 2B, Supplemental Fig. 2B**).

**Figure 2.**
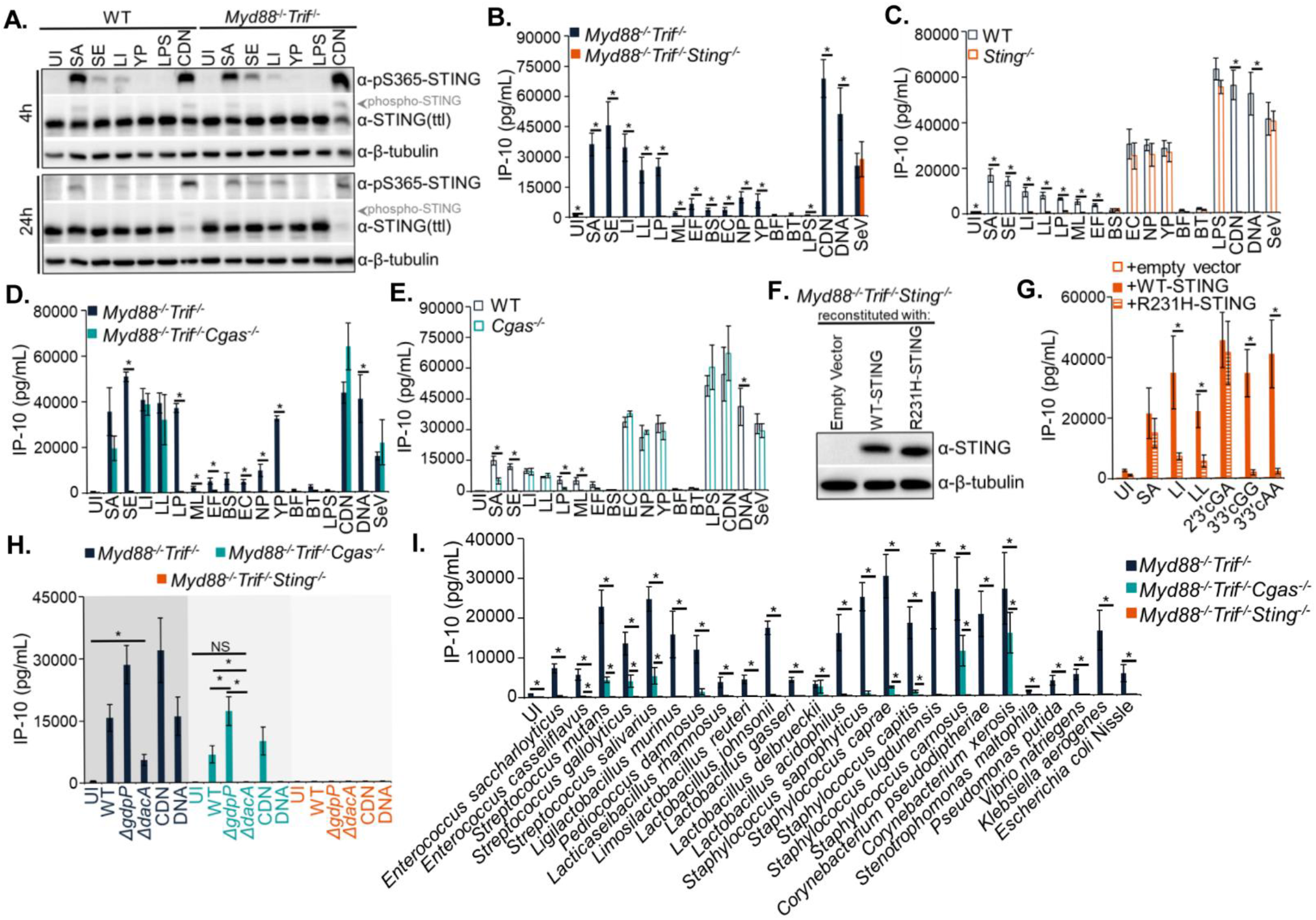
STING commonly detects diverse bacteria to promote IFN responses in macrophages. **(A)** WT and *Myd88*^-/-^*Trif*^-/-^ iBMDMs were treated as in **Fig. 1B**. 4 h (top) and 24 h (bottom) protein lysates were analyzed by immunoblot for phosphorylated STING (pS365), total STING, and p-Tubulin. **(B)** *Myd88*^-/-^*Trif*^-/-^ and *Myd88*^-/-^*Trif*^-/-^*Sting*^-/-^ iBMDMs were treated as in **Suppl. Fig. 1N**, with the additional treatment of cytosolically-delivered DNA (1 µg/mL). 24 h conditioned supernatants were analyzed by ELISA for IP-10. **(C)** WT and *Sting*^-/-^ iBMDMs were treated as in **Fig. 2B**, and 24 h conditioned supernatants were analyzed by ELISA for IP-10. **(D)** *Myd88*^-/-^*Trif*^-/-^ and *Myd88*^-/-^*Trif*^-/-^*Cgas*^-/-^ iBMDMs were treated as in **Fig. 2B**, and 24 h conditioned supernatants were analyzed by ELISA for IP-10. **(E)** WT and *Cgas*^-/-^ iBMDMs were treated as in **Fig. 2B**, and 24 h conditioned supernatants were analyzed by ELISA for IP-10. **(F)** *Myd88*^-/-^*Trif*^-/-^*Sting*^-/-^ iBMDMs were stably reconstituted with empty vector, WT-STING, or R231H-STING. Reconstituted cells were sorted for equal expression, as assessed by immunoblot for STING and β-tubulin. **(G)** Reconstituted **Fig. 2F** cells were exposed to bacteria as in **Fig. 1B**, in addition to 10 µg/mL CDNs. 24 h conditioned supernatants were analyzed by ELISA for IP-10. **(H)** *Myd88*^-/-^*Trif*^-/-^, *Myd88*^-/-^*Trif*^-/-^*Cgas*^-/-^, and *Myd88*^-/-^*Trif*^-/-^*Sting*^-/-^ iBMDMs were treated as in **Fig. 2B**. 24 h conditioned supernatants were analyzed by ELISA for IP-10. **(I)** *Myd88*^-/-^*Trif*^-/-^, *Myd88*^-/-^*Trif*^-/-^*Cgas*^-/-^, and *Myd88*^-/-^*Trif*^-/-^*Sting*^-/-^ iBMDMs were treated as in **Fig. 2B**. 24 h conditioned supernatants were analyzed by ELISA for IP-10.

These data demonstrate that STING is required for IFN responses to diverse bacteria.

To determine the interplay between STING and TLR signaling, we examined STING activation in WT cells where TLR signaling can be activated. We first generated *Sting*^*-/-*^ iBMDMs (**Supplemental Fig. 2C**). STING was required for IFNβ and IP-10 secretion in response to all gram-positive bacteria examined (**Fig. 2C, Supplemental Fig. 2D**). In contrast, WT and *Sting*^*-/-*^ iBMDMs secreted comparable amounts of IFNβ and IP-10 in response to gram-negative bacteria (**Fig. 2C, Supplemental Fig. 2D**). As we observed in WT iBMDMs, WT primary BMDMs exhibited increased STING activation upon exposure to bacteria (**Supplemental Fig. 2E**). The STING inhibitor H-151 (*24*) reduced IP-10 secretion from WT iBMDMs and primary BMDMs exposed to gram-positive, but not gram-negative, bacteria (**Supplemental Fig. 2F, G**). H-151 treatment of WT BMDMs also resulted in reduced STAT1 activation, as well as viperin production, only in response to gram-positive bacteria, as compared to vehicle-treated cells (**Supplemental Fig. 2H**). Genetic depletion or inhibition of STING resulted in modest reductions in TNFα secretion (**Supplemental Fig. 2I-K**). Bacterial survival in all genotypes was comparable (**Supplemental Fig. 2L**), and the absence of STING failed to alter bacterial association with cells (**Supplemental Fig. 2M, N**). STING and TLRs are therefore the principal mediators of IFN responses to bacteria in macrophages.

Because STING can detect CDNs produced by bacteria or cGAS, we examined the role for cGAS through generation of *Myd88*^*-/-*^*Trif*^*-/-*^*Cgas*^*-/-*^ iBMDMs (**Supplemental Fig. 2O**). As expected (*25*), *Myd88*^*-/-*^ *Trif*^*-/-*^*Cgas*^*-/-*^ iBMDMs were unable to secrete IFNβ and IP-10 upon transfection with DNA but responded normally to ADU-S100, which bypasses cGAS to activate STING directly (**Fig. 2D** and **Supplemental Fig. 2P**). *Myd88*^*-/-*^*Trif*^*-/-*^*Cgas*^*-/-*^ iBMDMs were defective for IFNβ and IP-10 secretion in response to several of the examined bacteria, as compared to *Myd88*^*-/-*^*Trif*^*-/-*^ iBMDMs (**Fig. 2D** and **Supplemental Fig. 2P**). However, cGAS was not required for IFN responses by *S. aureus, L. innocua*, or *L. lactis* (**Fig. 2D** and **Supplemental Fig. 2P**). cGAS-independent IFNβ and IP-10 responses to the same sets of bacteria were observed in iBMDMs singly deficient for cGAS (**Fig. 2E, Supplemental Fig. 2Q, R**).

Although cGAS is dispensable for IFN in response to *S. aureus, L. innocua*, or *L. lactis*, this finding does not preclude that some bacteria may have the ability to dually stimulate cGAS and STING during infection. To address this possibility, we utilized a STING allele containing an R231H mutation. R231H-STING responds to the cGAS-derived CDN, 2′3′-cGA, but is significantly less responsive to the bacteria-derived CDNs c-di-AMP (herein “3′3′-cAA”) and c-di-GMP (herein “3′3′-cGG”) (*26, 27*). These latter CDNs are known or predicted to be synthesized by *S. aureus, L. innocua*, and *L. lactis* (*28-30*). We reconstituted *Myd88*^*-/-*^*Trif*^*-/-*^*Sting*^*-/-*^ iBMDMs with WT- or R231H-STING, at equivalent protein abundance (**Fig. 2F**). As expected, WT- and R231H-STING enabled secretion of IP-10 in response to 2′3′-cGA, whereas WT-STING alone responded to 3′3′-cAA and 3′3′-cGG (**Fig. 2G**). We found that WT- and R231H-STING-producing cells secreted similar amounts of IP-10 in response to *S. aureus*. In contrast, we observed reduced IP-10 secretion to *L. innocua* and *L. lactis* in R231H-STING-expressing cells (**Fig. 2G**). Thus, while all bacteria examined in this assay stimulate STING directly, *S. aureus* also stimulates cGAS.

cGAS-independent stimulation of STING by bacteria implies that bacteria are the source for STING-stimulatory CDNs. To test this concept, we examined a *S. aureus* mutant lacking the diadenylate cyclase DacA, which is unable to synthesize 3′3′-cAA, or a mutant lacking the phosphodiesterase GdpP, which produces increased 3′3′-cAA levels (*31, 32*). In *Myd88*^*-/-*^*Trif*^*-/-*^*Cgas*^*-/-*^ iBMDMs, *ΔgdpP S. aureus* stimulated more IP-10 secretion compared to WT bacteria, and *ΔdacA S. aureus* failed to stimulate any IP-10 secretion (**Fig. 2H**). Thus, the abundance of IP-10 secretion correlates with 3′3′-cAA production in *S. aureus*. Consistent with our R231H-STING data that suggests *S. aureus* can also stimulate cGAS (**Fig. 2G**), *ΔdacA S. aureus* stimulated IP-10 secretion from *Myd88*^*-/-*^*Trif*^*-/-*^ iBMDMs, where cGAS is intact (**Fig. 2H**). These data suggest that the production of CDNs by bacteria is required for direct STING stimulation but dispensable for cGAS activation.

Taken together, our use of evolutionarily diverse bacteria for side-by-side macrophage infections revealed a common role for cGAS and/or STING in bacterial detection. To test the universality of these findings, we next examined a larger panel of 25 evolutionarily diverse bacteria (**Supplemental Fig. 2S)**. Consistent with our above findings, all of these diverse gram-positive and gram-negative bacteria we examined stimulated IP-10 secretion in WT and *Myd88*^*-/-*^*Trif*^*-/-*^ iBMDMs (**Fig. 2I** and **Supplemental Fig. 2T**). Loss of STING in *Myd88*^*-/-*^*Trif*^*-/-*^ iBMDMs resulted in a complete loss of IP-10 secretion to all bacteria tested (**Fig. 2I**). cGAS accounted for detection of only a subset of bacteria as several of the examined bacteria still stimulated IP-10 secretion in *Myd88*^*-/-*^*Trif*^*-/-*^*Cgas*^*-/-*^ iBMDMs (**Fig. 2I**). As demonstrated with our earlier panel of bacteria, this reliance on cGAS and/or STING for IP-10 secretion persisted in otherwise WT iBMDMs for gram-positive, but not for gram-negative, bacteria (**Supplemental Fig. 2T**).

### STING detection of bacteria occurs irrespective of virulence potential

Many of the STING-stimulating bacteria in our panel are associated with humans and rarely cause disease (summarized in **Supplemental Table 1**) (*33-44*), yet pathogenic bacteria and their membrane-damaging virulence factors have been most associated with activation of cytoplasmic PRRs (*5, 6*). We therefore determined the role of virulence in STING activation by bacteria. We first assessed STING activities during infections with the pathogen *Listeria monocytogenes*. The cytolysin, Listeriolysin O (encoded by the *hly* gene), along with two phospholipases, PlcA and PlcB, are required for *L. monocytogenes* to damage and escape phagosomes (*45*). Several studies concluded that escape from phagosomes is a prerequisite for STING-dependent IFN responses to *L. monocytogenes* (*45-50*). We examined these prior findings through the study of WT and *ΔhlyΔplcAΔplcB L. monocytogenes* and the related non-pathogen *L. innocua*, which does not encode for *hly* or escape from phagosomes (*51*). At 4 hours post-infection, only WT *L. monocytogenes*, but not the *ΔhlyΔplcAΔplcB* mutant or *L. innocua*, stimulated STAT1 activation and viperin production by iBMDMs (**Fig. 3A**). Interestingly, by extending the infection to 24 hours, all *Listeria* strains induced IFN responses, as assessed by cellular abundance of pSTAT1, total STAT1, and viperin and secretion of IP-10 from *Myd88*^*-/-*^*Trif*^*-/-*^ iBMDMs (**Fig. 3A, B**). A higher MOI was needed for *ΔhlyΔplcAΔplcB* and *L. innocua* to generate an IFN response magnitude comparable to WT *L. monocytogenes* (**Fig. 3A, B**). IP-10 secretion in response to all listeriae required STING, but not cGAS (**Fig. 3B**).

**Figure 3.**
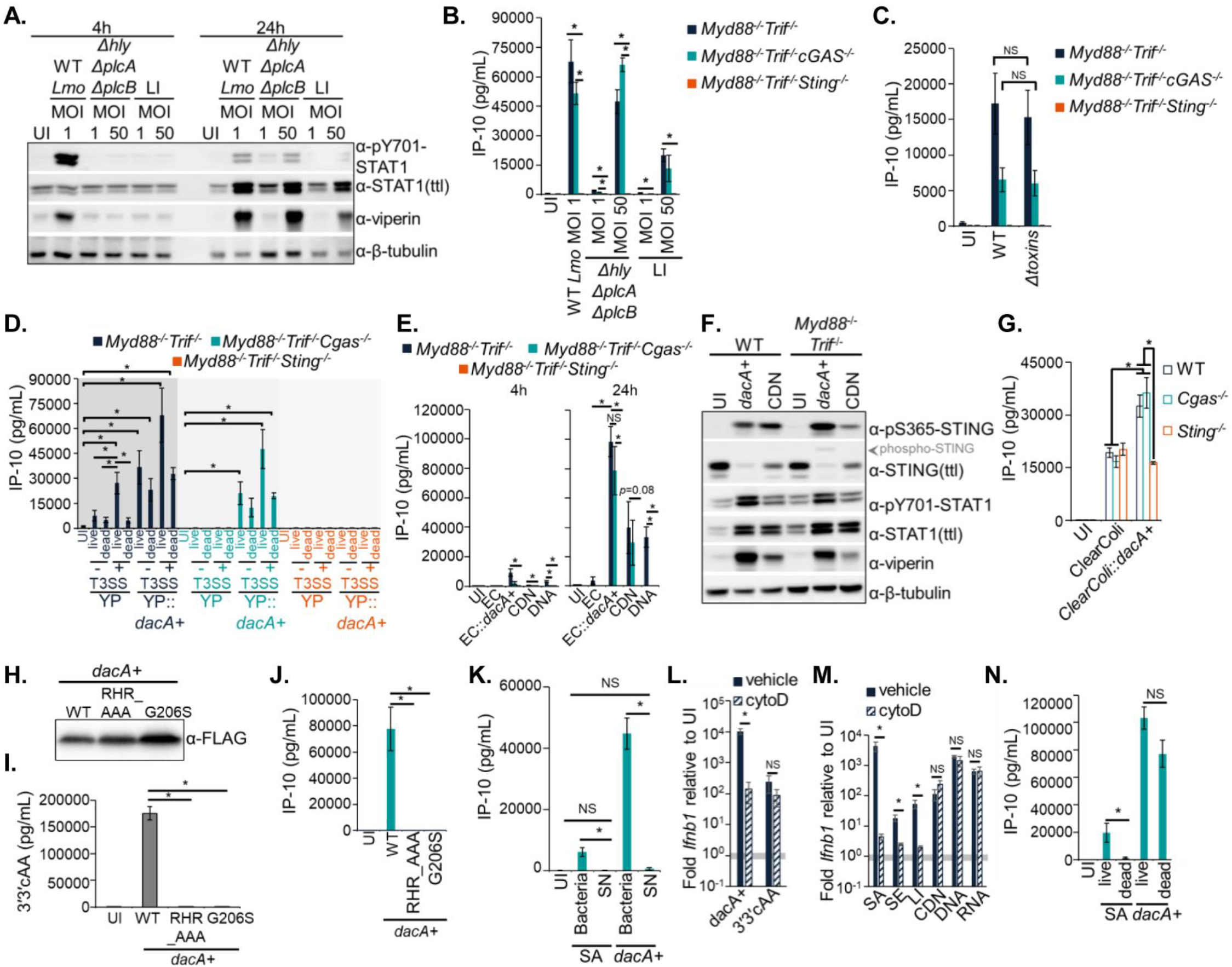
STING detects bacteria irrespective of virulence. **(A)** *Myd88*^-/-^*Trif*^-/-^ iBMDMs were exposed to WT *L. monocytogenes* (“Lmo”), *ΔhlyΔpIcAΔpIcB Lmo*, or *L. innocua* (“LI”) at the indicated MOIs. 4 h (left) and 24 h (right) protein lysates were analyzed by immunoblot for pSTATI, total STAT1, viperin, and β-tubulin. **(B)** *Myd88*^-/-^*Trif*^-/-^, *Myd88*^-/-^*Trif*^-/-^*Cgas*^*-/-*^ and *Myd88*^-/-^*Trif*^-/-^*Sting*^*-/-*^ iBMDMs were treated as in **Fig. 3A**. 24 h conditioned supernatants were analyzed by ELISA for IP-10. **(C)** *Myd88*^-/-^*Trif*^-/-^*Myd88*^-/-^*Trif*^-/-^*Cgas*^*-/-*^ and *Myd88*^-/-^*Trif*^-/-^*Sting*^*-/-*^ iBMDMs were treated as in **Fig. 1B**. 24 h conditioned supernatants were analyzed by ELISA for IP-10. **(D)** *Y. pseudotuberculosis* (“YP”) and *Y. pseudotuberculosis::dacA*+ (“YP::*dacA+*”) were grown in type 3 secretion system-inducing conditions (“T3SS+”) or not (“T3SS-”), with IPTG (1mM) to induce production of DacA-1xFLAG. Bacteria were then untreated (“live”) or treated with paraformaldehyde for 15 min (“dead”). *Myd88*^-/-^*Trif*^-/-^,*Myd88*^-/-^*Trif*^-/-^*Cgas*^*-/-*^, and *Myd88*^-/-^*Trif*^-/-^*Sting*^*-/-*^ iBMDMs were exposed to bacteria at an MOI of 50. 24 h conditioned supernatants were analyzed by ELISA for IP-10. **(E)** *E. coli* (“EC”) and *E. coli::dacA*+ (“EC*::dacA*+”) were grown with IPTG (1 mM) to induce production of DacA-3xFLAG. *Myd88*^-/-^*Trif*^-/-^, *Myd88*^-/-^*Trif*^-/-^*Cgas*^*-/-*^, and *Myd88*^-/-^*Trif*^-/-^*Sting*^*-/-*^ iBMDMs were treated as in **Fig. 1B**. 24 h conditioned supernatants were analyzed by ELISA for IP-10. **(F)** WT and *Myd88*^-/-^*Trif*^-/-^ iBMDMs were treated as in **Fig. 1B** and IPTG (0.01 mM)-treated *E. coli::dacA+* at an MOI of 50. 24 h protein lysates were analyzed by immunoblot for pSTING, total STING, pSTATI, total STAT1, viperin, and β-tubulin. **(G)** WT, *Cgas*^*-/-*^, and *Sting*^*-/-*^ iBMDMs were treated as in **Fig. 1B** and IPTG (0.01 mM)-treated *ClearColi* and *ClearColi::dacA*+ at an MOI of 50. 24 h conditioned supernatants were analyzed by ELISA for IP-10. **(H)** *E. coli* were treated with IPTG (1 mM) to induce expression of WT-DacA-3xFLAG, RHR_AAA-DacA-3xFLAG, or G206S-DacA-3xFLAG. Protein lysates were collected from bacteria and analyzed by immunoblot for FLAG protein. **(I)** Bacteria from **Fig. 3H** were lysed and analyzed for the presence of 3’3’cAA by ELISA. **(J)** *Myd88*^-/-^ *Trif*^-/-^ *Cgas*^-/-^ iBMDMs exposed to bacteria from **Fig. 3H** at an MOI of 50. 24 h conditioned supernatants were analyzed by ELISA for IP-10. **(K)** Bacteria and conditioned supernatants (“SN”) were prepared from 3h S. aureus and E. coli::dacA+ mid-log cultures. *Myd88*^-/-^ *Trif*^-/-^*Cgas*^-/-^ iBMDMs were exposed to bacteria at an MOI of 50 or 1 mL of SN (see methods). 24 h infection conditioned supernatants were analyzed by ELISA for IP-10. **(L)** *Myd88*^-/-^ *Trif*^-/-^ iBMDMs were treated with cytochalasin D (“cytoD”) or not (“vehicle”) prior to and throughout exposure to IPTG (0.01 mM)-treated E. coli::dacA+ at an MOI of 50 and 3’3’cAA (10 μg/mL). 6 h RNA lysates were analyzed for fold Ifnbl transcriptional responses as in **Fig. 1C**. **(M)** *Myd88*^-/-^ *Trif*^-/-^ iBMDMs were treated with cytochalasin D as in **Fig. 3L** and bacteria and stimuli as in **Fig. 2B**, with the additional stimuli of cytosolically-delivered 5’ppp-dsRNA (“RNA”; 0.5 μg/mL). 6 h RNA lysates were analyzed for fold Ifnbl transcriptional responses as in **Fig. 1C**. **(N)** *Myd88*^-/-^ *Trif*^-/-^ *Cgas*^-/-^ iBMDMs were exposed to live and dead S. aureus and IPTG (ImM)-treated E. coli::dacA+ (see methods) at an MOI of 50. 24 h conditioned supernatants were analyzed by ELISA for IP-10.

*S. aureus* produces a number of pore-forming toxins that are critical to its virulence and activation of cytosolic sensors that seed assembly of inflammasomes (*52, 53*). To test whether pore-forming toxins mediate STING activation to *S. aureus*, we utilized a *Δtoxins* strain lacking all major pore-forming toxins (*54*). Parental and *Δtoxins S. aureus* similarly stimulated IP-10 secretion in *Myd88*^*-/-*^*Trif*^*-/-*^ and *Myd88*^*-/-*^*Trif*^*-/-*^*Cgas*^*-/-*^ iBMDMs (**Fig. 3C**). Thus, the major pore-forming toxins of *S. aureus* are dispensable for STING detection of these bacteria.

A similar set of studies was performed with a strain of *Y. pseudotuberculosis*, which encodes all of the components of a membrane-damaging type 3 secretion system (T3SS) apparatus but lacks the effectors that modulate immune signaling (*55-57*). We engineered this *Y. pseudotuberculosis* strain to produce *S. aureus* DacA to synthesize 3′3′-cAA (**Supplemental Fig. 3A**). The parental strain of *Y. pseudotuberculosis* stimulated IP-10 secretion from *Myd88*^*-/-*^*Trif*^*-/-*^ iBMDMs, by a process that required expression of the T3SS, bacterial viability, cGAS and STING (**Fig. 3D**). In contrast, *Y. pseudotuberculosis::dacA+* stimulated IP-10 secretion in a cGAS-independent (but STING-dependent) manner, regardless of T3SS expression or bacterial viability (**Fig. 3D**). These findings using gram-positive (*Listeria* and *S. aureus*) and gram-negative (*Yersinia*) bacteria support the concept that virulence, in this case via expression of membrane-damaging proteins, can potentiate STING activation but is not strictly required for STING activation.

Encouraged by the magnitude of cGAS-independent IP-10 secretion to *Y. pseudotuberculosis::dacA+* (**Fig. 3D**), we sought to further define the properties underlying bacteria-mediated STING stimulation by engineering a simple, non-pathogenic laboratory strain of *E. coli* to produce *S. aureus* DacA (**Supplemental Fig. 3B**). Parent *E. coli* stimulated modest IP-10 secretion from *Myd88*^*-/-*^ *Trif*^*-/-*^ iBMDMs in a manner requiring cGAS and STING (**Fig. 3E**). In contrast, *E. coli::dacA+* stimulated robust IP-10 secretion in *Myd88*^*-/-*^*Trif*^*-/-*^ iBMDMs in a manner independent of cGAS but requiring STING (**Fig. 3E**). WT and *Myd88*^*-/-*^*Trif*^*-/-*^ iBMDMs exposed to *E. coli::dacA+* exhibited increased detection of pSTING, pSTAT1, and viperin, compared to uninfected cells, and at this late time point of 24 hours, total STING protein abundance was reduced upon exposure to *E. coli::dacA+* or ADU-S100 in WT and *Myd88*^*-/-*^*Trif*^*-/-*^ iBMDMs, compared with uninfected cells (**Fig. 3F**). This decrease in STING abundance is expected as STING is degraded as a negative regulatory mechanism (*3*). Notably, our infections with natural bacteria did not result in obvious STING degradation under similar conditions (**Fig. 2A**), suggesting that a larger proportion of total STING protein is activated by engineered *E. coli* producing CDNs. DacA production in a strain of *E. coli* whose LPS poorly stimulates TLR4 (**Supplemental Fig. 3C**, see methods) (*58*) significantly increased IP-10 secretion from WT iBMDMs, compared to parental bacteria (**Fig. 3G**). Increased IP-10 secretion required STING, but not cGAS, in this context (**Fig. 3G**). Thus, heterologous production of DacA is sufficient to elicit robust IFN responses and to convert non-pathogenic *E. coli* from a direct agonist of cGAS to a direct STING agonist.

To manipulate 3′3′-cAA levels during infections with *E. coli::dacA+*, we first decreased DacA production by altering the concentration of IPTG used for expression of *dacA* in *E. coli* (**Supplemental Fig. 3D**). Decreasing IPTG concentrations resulted in a step-wise decrease in IP-10 secretion from *Myd88*^*-/-*^ *Trif*^*-/-*^*Cgas*^*-/-*^ iBMDMs in response to *E. coli::dacA+* (**Supplemental Fig. 3E**). Similarly, decreasing the MOI diminished the extent of IP-10 secretion from these cells (**Supplemental Fig. 3F**). An MOI as low as 0.1 still induced robust IP-10 production (∼25 ng from ∼10^6^ cells) (**Supplemental Fig. 3F**). To determine whether DacA enzymatic activity is important for cGAS-independent IFN responses, we generated *E. coli* expressing catalytically inactive *dacA* mutants where the codons for R208, H209, and R210 were swapped for codons encoding alanine or where the codon for G206 was swapped for a codon encoding serine (*59, 60*). DacA mutants were equivalently abundant in *E. coli* (**Fig. 3H**), yet 3′3′-cAA was only detected from lysed *E. coli* producing WT-DacA (**Fig. 3I**). Only *E. coli* producing WT-DacA stimulated IP-10 secretion from *Myd88*^*-/-*^*Trif*^*-/-*^*Cgas*^*-/-*^ iBMDMs (**Fig. 3J**). DacA enzymatic activity and protein abundance therefore define 3′3′-cAA levels and the subsequent magnitude of direct STING stimulation.

As pure CDNs added exogenously to macrophages can enter cells and stimulate STING (*16, 17*), we explored the possibility that 3′3′-cAA was released from extracellular *E. coli::dacA+* to activate IFN responses. To test this possibility, we harvested conditioned supernatants from *S. aureus* and *E. coli::dacA+* cultures, which both synthesize 3′3′-cAA to stimulate STING. While live *S. aureus* and *E. coli::dacA+* stimulated IP-10 secretion from *Myd88*^*-/-*^ *Trif*^*-/-*^*Cgas*^*-/-*^ iBMDMs, their conditioned supernatants failed to induce IP-10 secretion from these cells (**Fig. 3K**). This finding suggests that intracellular *E. coli::dacA+* bacteria are the source of CDNs that stimulate STING. Consistent with this idea, preventing phagocytosis of *E. coli::dacA+* by cytochalasin D treatment inhibited *Ifnb1* transcription (**Fig. 3L**). To determine a general role of phagocytosis, we examined the impact of cytochalasin D on IFN responses to *S. aureus, S. epidermidis*, and *L. innocua*, compared with pure ligands for STING, cGAS, and the RLR RIG-I. Cytochalasin D treatments specifically prevented *Ifnb1* expression by macrophages exposed to bacteria (**Fig. 3M**). Thus, phagocytosis is a general prerequisite for STING activation in response to natural and bioengineered bacteria.

We considered the possibility that *E. coli* actively promote STING activation through the damage or manipulation of phagosomes. However, live and dead *E. coli::dacA+* stimulated comparable secretion of IP-10 from *Myd88*^*-/-*^*Trif*^*-/-*^*Cgas*^*-/-*^ iBMDMs (**Fig. 3N**). Additionally, we found *E. coli::dacA+* survived poorly within macrophages (**Supplemental Fig. 3G**). These findings suggest that release of CDNs from dead bacteria within phagosomes is a means to stimulate STING-dependent IFN responses.

### The CDN transporters SLC46A2 and LRRC8A mediate STING activation by bacteria

We hypothesized that in lieu of bacterial manipulation or damage of phagosomes, host factors mediate CDN passage across the phagosomal membrane. Several membrane-spanning transport proteins can facilitate entry of pure CDNs into the cytosol (*61*). These include members of the solute carrier (SLC) family of transporters (*i*.*e*., SLC19A1, SLC46A1, SLC46A2, and SLC46A3), as well as the volume-regulated anion channel (VRAC) family, which are heterohexameric channels composed of an obligatory protein, termed leucine-rich repeat-containing protein 8A (LRRC8A) (*16-20*). All of these proteins are thought to function in CDN transport from the plasma membrane, yet their activities have only been documented using pure CDNs (*16-20*). Curiously, SLC46A1, SLC46A2, SLC46A3, LRRC8A, and fluorescently-labelled CDNs can all localize to the endosomal network, and endocytosis enhances cytosolic entry of CDNs (*14, 62-65*). We therefore considered the potential role for these CDN transport proteins in STING activation by phagosomal bacteria. We first examined a role for SLCs. mRNA transcripts for each SLC transporter were detected in WT and *Myd88*^*-/-*^*Trif*^*-/-*^ iBMDMs, except for *Slc46a2*, as previously noted (**Supplemental Fig. 4A**) (*18*). We generated *Myd88*^*-/-*^*Trif*^*-/-*^ iBMDMs deficient for *Slc19a1, Slc46a1, Slc46a2*, or *Slc46a3*. None of these SLC deficiencies impacted IP-10 secretion in response to 2′3′-cGA or CDN-producing bacteria (**Supplemental Fig. 4B-E**). We next stably overexpressed SLCs 19A1, 46A1, 46A2, or 46A3 in *Myd88*^*-/-*^*Trif*^*-/-*^ iBMDMs (**Supplemental Fig. 4F**). Overexpression of *Slc46a2*, but not other SLCs, increased IP-10 secretion induced by the pure CDN 2′3′-cGA and the CDN-producing bacteria *E. coli::dacA+, L. innocua*, and *L. lactis* (**Fig. 4A, B, Supplemental Fig. 4G-I**). Increased IP-10 correlated with increased pSTING and viperin protein abundance to *E. coli::dacA+* in *Slc46a2*-overexpressing cells (**Fig. 4C**). *Slc46a2* overexpression did not impact IP-10 secretion in response to the membrane-permeant STING agonist DMXAA (**Fig. 4A, B**). Thus, SLC46A2 is sufficient to mediate IFN responses to CDN-producing bacteria.

**Figure 4.**
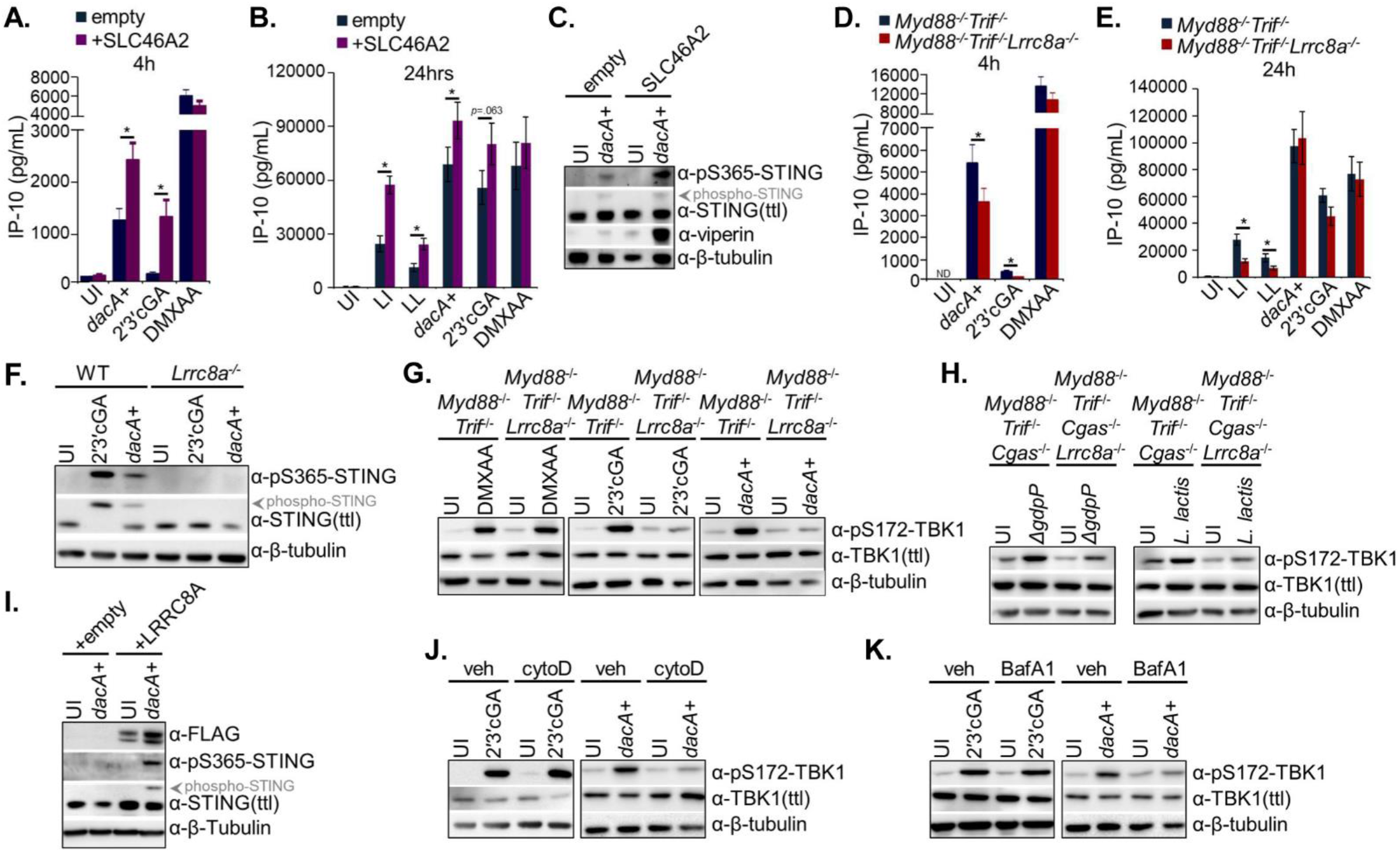
The CDN transporters LRRC8A and SLC46A2 support bacteria-driven STING activation. **(A)** *Myd88*^-/-^ *Trif*^-/-^ iBMDMs reconstituted with empty vector or SLC46A2-3xFLAG were treated as in **Suppl. Fig. 4G**. 4 h conditioned supernatants were analyzed by ELISA for IP-I0. **(B)** *Myd88*^-/-^ *Trif*^-/-^ iBMDMs reconstituted with empty vector or SLC46A2-3xFLAG were treated as in **Suppl. Fig. 4G**. 24 h conditioned supernatants were analyzed by ELISA for IP-I0. **(C)** Empty vector or *SIc46a2-overexpressing Myd88*^-/-^ *Trif*^-/-^ *Lrrc8a*^-/-^ iBMDMs were treated as in **Suppl. Fig. 4G**. 4 h protein lysates were analyzed by immunoblot for pSTlNG, total STING, viperin, and β-tubulin. **(D)** *Myd88*^-/-^ *Trif*^-/-^ and *Myd88*^-/-^ *Trif*^-/-^ *Lrrc8a*^-/-^ iBMDMs were treated as in **Suppl. Fig. 4G**. 4 h conditioned supernatants were analysed by ELISA for IP-I0. **(E)** *Myd88*^-/-^ *Trif*^-/-^ and *Myd88*^-/-^ *Trif*^-/-^ *Lrrc8a*^-/-^ iBMDMs were treated as in **Suppl. Fig. 4G**. 24 h conditioned supernatants were analyzed by ELISA for IP-I0. **(F)** WT and *Lrrc8a*^-/-^ iBMDMs were treated as in **Suppl. Fig. 4N**. 4 h protein lysates were analyzed by immunoblot for pSTlNG, total STING, pTBK1, total TBK1, and β-tubulin. **(G)** *Myd88*^-/-^ *Trif*^-/-^ and *Myd88*^-/-^ *Trif*^-/-^ *Lrrc8a*^-/-^ iBMDMs were treated as in **Suppl. Fig. 4G** under physiologic conditions for 1 h. Cells were then chased with hypotonic conditions for 4 h. 5 h protein lysates were analyzed by immunoblot for pTBK1, total TBK1, and β-tubulin. **(H)** *Myd88*^-/-^ *Trif*^-/-^ *Cgas*^-/-^and *Myd88*^-/-^ *Trif*^-/-^ *Cgas*^-/-^ *Lrrc8a*^-/-^ iBMDMs were exposed to *ΔgdpP S. aureus* and *L. lactis* at an MOI of 50 under physiologic conditions for 4.5 h. Cells were then chased with hypotonic conditions for 1.5 h. 6h protein lysates were analyzed by immunoblot for pTBK1, total TBK1, and β-tubulin, **(I)** *Lrrc8a*^-/-^ iBMDMs stably reconstituted with empty vector or a doxycycline-inducible LRRC8A-3xFLAG were exposed as in **Suppl. Fig. 4G** under physiologic conditions for 1 h. Cells were then chased with hypotonic conditions for 3h. 4 h protein lysates were analyzed by immunoblot for FLAG, pSTlNG, total STING, and β-tubulin. **(J)** *Myd88*^-/-^ *Trif*^-/-^ iBMDMs were treated with cytochalasin D as in **Fig. 3L** and treated with the stimuli as in **Suppl. Fig. 4G** under hypotonic conditions. 4 h protein lysates were analyzed by immunoblot for pTBK1, total TBK1, and β-tubulin. **(K)** *Myd88*^-/-^ *Trif*^-/-^ iBMDMs were treated in the absence (“veh”) or presence of bafilomycin A1 (“BafA1 “) prior to or throughout exposure to stimuli in **Suppl. Fig. 4G**. 4 h protein lysates were analyzed by immunoblot for pTBK1, total TBK1, and β-tubulin.

As genetic deficiency in the above SLCs failed to affect IFN responses to CDNs or bacteria, we examined a role for VRACs. mRNA transcripts for all five LRRC8 components of VRACs were detectable in WT and *Myd88*^*-/-*^*Trif*^*-/-*^ iBMDMs (**Supplemental Fig. 4A**). As LRRC8A is an obligatory component of each VRAC channel, we generated *Myd88*^*-/-*^*Trif*^*-/-*^*Lrrc8a*^*-/-*^ iBMDMs (**Supplemental Fig. 4J**). At all examined time points, IP-10 secretion to the membrane permeable STING agonist DMXAA was similar between *Myd88*^*-/-*^ *Trif*^*-/-*^ and *Myd88*^*-/-*^*Trif*^*-/-*^*Lrrc8a*^*-/-*^ iBMDMs (**Fig. 4D, E**). In contrast, IP-10 secretion to 2′3′-cGA at 4 hours was reduced in *Myd88*^*-/-*^*Trif*^*-/-*^*Lrrc8a*^*-/-*^ iBMDMs, compared to *Myd88*^*-/-*^*Trif*^*-/-*^ cells (**Fig. 4D, E**). The partial defect in IFN responses to pure 2′3′-cGA in *Lrrc8a*-deficient macrophages is consistent with previous reports (*19, 20*). We observed that IP-10 secretion was also reduced in *Myd88*^*-/-*^*Trif*^*-/-*^*Lrrc8a*^*-/-*^ iBMDMs to *E. coli::dacA+* and the natural CDN-producing bacteria *L. innocua* and *L. lactis* (**Fig. 4D, E**). These data suggest that LRRC8A mediates IFN responses to CDN-producing bacteria.

As VRAC transport activities are enhanced under hypotonic conditions (*19, 20*), we speculated that the phenotype associated with *Lrrc8a*-deficiency and CDN-producing bacteria may be enhanced when cells are cultured in hypotonic media. We first examined STING activation in *Myd88*^*-/-*^*Trif*^*-/-*^ and *Myd88*^*-/-*^*Trif*^*-/-*^*Lrrc8a*^*-/-*^ iBMDMs cultured under physiologic and hypotonic medium. Regardless of culture condition, *Lrrc8a*-deficiency did not impact pSTING abundance induced by transfection of cells with DNA (**Supplemental Fig. 4K**). Under physiologic conditions, *Lrrc8a*-deficiency did not impact STING activation within cells exposed to exogenous 2′3′-cGA (**Supplemental Fig. 4K, L**). Hypotonic conditions, in contrast, enhanced the ability of 2′3′-cGA to activate STING and TBK1, by a process that required LRRC8A (**Supplemental Fig. 4K, L**). Thus, consistent with previous work (*19, 20*), LRRC8A is required for STING activation induced by pure, exogenous 2′3′-cGA under channel-gating conditions.

LRRC8A was similarly required for STING and TBK1 activation in response to *E. coli::dacA+* under channel-gating hypotonic conditions (**Fig. 4F, Supplemental Fig. 4M, N**). This finding was made with two different *Myd88*^*-/-*^*Trif*^*-/-*^*Lrrc8a*^*-/-*^ clones generated from two different CRISPR guides, as well as *Lrrc8a*-deficiency on *Myd88*^*-/-*^*Trif*^*-/-*^*Cgas*^*-/-*^ and WT backgrounds (**Supplemental Fig. 4J**). Inhibition of VRAC channels with the chemical DCPIB also reduced TBK1 activation by *E. coli::dacA+* in *Myd88*^*-/-*^*Trif*^*-/-*^ iBMDMs under hypotonic conditions (**Supplemental Fig. 4O**). Under our hypotonic conditions, all iBMDM genotypes maintained ∼90% viability (**Supplemental Fig. 4P**). Notably, LRRC8A did not impact the phagocytic capacity of cells (**Supplemental Fig. 4Q, R**). This finding is consistent with the idea that LRRC8A mediates STING activation to bacteria downstream of phagocytosis. To further mitigate any off-target effect of hypotonicity on macrophage functions, we performed pulse-chase experiments where phagocytosis occurred in physiologic media, followed by a short treatment of cells with hypotonic media. In the extreme example of this pulse-chase regimen, a one hour treatment with hypotonic media after three hours of host-bacteria interactions in physiological media was sufficient to reveal a requirement for LRRC8A for IFN responses to *E. coli::dacA+* (**Supplemental Fig. 4S**). Similar findings were made with 2′3′-cGA, *L. lactis*, and the 3′3′-cAA-overproducing *ΔgdpP S. aureus* mutant (**Fig. 4G, H**). In contrast, DMXAA-induced IFN responses were not sensitive to LRRC8A deficiency (**Fig. 4G**). To confirm the specificity of *Lrrc8a* depletion on these phenotypes, we reconstituted *Lrrc8a*^*-/-*^ iBMDMs with C-terminally-3xFLAG-tagged LRRC8A under a doxycycline-inducible promoter. *Lrrc8a*^*-/-*^ iBMDMs reconstituted with *Lrrc8a*, but not empty vector, restored STING activation in response to *E. coli::dacA+* under pulse-chase conditions (**Fig. 4I**). We found that genetic deficiency in *Lrrc8a*, but not other *Lrrc8* genes, resulted in defective TBK1 activation in response to *E. coli::dacA+* and 2′3′-cGA (**Supplemental Fig. 4T**).

We next examined how phagocytosis and the phagosome contributes to STING detection of CDN-producing bacteria under hypotonic conditions. Treatment with cytochalasin D reduced TBK1 activation in response to *E. coli::dacA+*, but not 2′3′-cGA, in *Myd88*^*-/-*^*Trif*^*-/-*^ iBMDMs under hypotonic conditions (**Fig. 4J**). We then determined if the acidified, bacteriolytic environment of the phagosome is required for bacteria-induced STING activation. Cells treated with bafilomycin A1, an inhibitor of phagosome acidification, prevented TBK1 activation in response to *E. coli::dacA+*, but not 2′3′-cGA, in *Myd88*^*-/-*^*Trif*^*-/-*^ iBMDMs under hypotonic conditions (**Fig. 4K**). These findings inform a model whereby CDN-producing bacteria are first internalized into bacteriolytic phagosomes, followed by STING activation mediated by the CDN transporters LRRC8A or SLC46A2.

To define the relationship between LRRC8A and SLC46A2, we first overexpressed *Slc46a2* in *Lrrc8a*^*-/-*^ iBMDMs (**Supplemental Fig. 4U**). *Lrrc8a*^*-/-*^ iBMDMs overexpressing *Slc46a2*, but not empty vector, exhibited increased STING activation in response to *E. coli::dacA+* (**Supplemental Fig. 4V**). SLC46A2 therefore operates independently of LRRC8A. We next examined TBK1 activation in SLC-deficient cells exposed to *E. coli::dacA+* under LRRC8A-gating hypotonic conditions. Only *Myd88*^*-/-*^ *Trif*^*-/-*^*Lrrc8a*^*-/-*^ iBMDMs exhibited defects in TBK1 activation in response to *E. coli::dacA+* (**Supplemental Fig. 4W**). To address the possibility that SLCs may functionally compensate for the loss of LRRC8A, we generated *Myd88*^*-/-*^*Trif*^*-/-*^*Lrrc8a*^*-/-*^*Slc19a1*^*-/-*^*Slc46a1*^*-/-*^*Slc46a2*^*-/-*^*Slc46a3*^*-/-*^ iBMDMs. However, this combinatorial genetic depletion failed to alter IP-10 secretion, compared with *Myd88*^*-/-*^*Trif*^*-/-*^*Lrrc8a*^*-/-*^ iBMDMs (**Supplemental Fig. 4X**). A context-dependent network of CDN transporters may therefore function within cells to link phagosomal bacteriolysis to STING activation. This finding is akin to observations with other small molecule transporters, such as in the case of pH-regulation where no fewer than 63 human H^+^ transporters function from the plasma membrane alone (*66*).

## Discussion

Our findings highlight STING as a versatile PRR that rivals the TLRs in its ability to detect a wide range of bacterial encounters. Our understanding for the general principles underlying cellular detection of bacteria have been hindered by a historical focus on disease-causing bacteria (*i*.*e*., opportunistic or strict pathogens), even though the host’s most common encounters are with bacteria in the absence of disease (*i*.*e*., non-pathogens). STING is no exception to this pathogen-oriented research bias, with the majority of studies examining pathogens, revealing both host-protective and host-maladaptive roles for STING in the control of pathogenic infections (*67*). These studies also implicated pathogen-mediated membrane damage as a determinant for STING activation (*10*). Our use of diverse bacteria tested this pathogen-specific view, and revealed STING, but not other unrelated cytoplasmic PRRs, as a generic sensor of bacteria to drive IFN responses. This work also highlights the complexity and overlapping pathways (*i*.*e*., TLRs, cGAS, and STING) that ensures IFN as a common outcome of macrophage-bacteria interactions. Our findings mirror those made *in vivo* where commensal bacteria drive basal IFN in phagocytes in a manner requiring multiple PRRs, including STING and TLRs (*68-77*).

The versatility of STING as a generic sensor of bacteria relies on the cytosolic delivery of their CDNs (for direct STING agonism) and, potentially, their DNA (for cGAS agonism). The cytosolic delivery of CDNs, in particular, can only be partially explained by membrane-perturbing virulence factors as these factors were either entirely dispensable or only partially required for STING activation. Moreover, we discovered several non-pathogenic bacteria that could stimulate STING directly, including *L. innocua, L. lactis, Corynebacterium xerosis*, several commensal Streptococci, and several commensal Staphylococci. Our findings suggest that the plasma membrane is not the primary conduit taken by bacteria-produced CDNs to access the cytosol. Rather, this process occurs after phagocytosis and bacteriolysis, leading to SLC46A2- and LRRC8A-mediated translocation of CDNs to STING. As most sequenced bacteria encode at least one CDN synthase (*78, 79*), the findings reported herein provide a mandate to explore the potential roles for STING in host-bacteria interactions, regardless of virulence potential, and to explore the interplay and *in vivo* immunological consequences between bacteria-driven activation of cGAS compared to direct STING stimulation.

## Materials and Methods

### Phylogenetic Analysis

Bacterial species and their 16S ribosomal RNA sequences (NCBI reference sequence number in parentheses below) were collated from 6 of the most common human-associated phyla. Bacteria in asterisks (*) were used as references for bacteria in the **Fig. 1** (^1^) or **Supplemental Fig. 2** trees (^2^). \**Erysipelothrix rhusiopathiae* (NR_040837.1), \**Parvimonas micra* (NR_036934.1), ^1^*L. innocua* (NR_029318.1), ^1^*B. subtilis* (NR_112116.2), ^1^*S. aureus* (NR_118997.2), ^1^*S. epidermidis* (NR_036904.1), ^1^*L. lactis* (NR_040955.1), ^1^*L. plantarum* (MG066537.1), ^1^*E. faecalis* (NR_040789.1), \**Clostridium perfringens* (NR_121697.2), ^2^*Staphylococcus carnosus* (NR_027518.1), ^2^*Staphylococcus capitis* (NR_027519.1), ^*2*^*Staphylococcus caprae* (NR_024665.1), ^*2*^*Staphylococcus lugdunensis* (NR_024668.1), ^*2*^*Staphylococcus saprophyticus* (NR_074999.2), ^*2*^*Lactobacillus acidophilus* (NR_043182.1), ^*2*^*Lactobacillus delbrueckii* (NR_029106.1), ^*2*^*Lacticaseibacillus rhamnosus* (NR_043408.1), ^*2*^*Limosilactobacillus reuteri* (NR_025911.1), ^*2*^*Lactobacillus gasseri* (NR_075051.2), ^*2*^ *Ligilactobacillus murinus* (NR_042231.1), ^*2*^*Lactobacillus johnsonii* (NR_025273.1), ^*2*^*Enterococcus casseliflavus* (NR_104560.1), ^*2*^*Enterococcus Streptococcus* (NR_041707.1), ^*2*^ *Streptococcus salivarius* (NR_042776.1), ^*2*^ *Streptococcus mutans*(NR_042772.1), ^*2*^ *Streptococcus gallolyticus* (NR_044904.1) ^*2*^*Pediococcus damnosus* (NR_042087.1), and *Dialister *invisus* (NR_025680.1) were used as Firmicutes representatives. ^1^*B. fragilis* (NR_074784.2), ^1^*B. theta* (NR_074277.1), \**Chitinophaga terrae* (NR_041540.1), \**Elizabethkingia meningoseptica* (NR_042267.1), and \**Sphingobacterium spiritivorum* (NR_044077.1) were used as Bacteroidetes representatives. \**Corynebacterium glucuronolyticum* (NR_036792.1), ^*2*^*Corynebacterium pseudodiphtheriticum* (NR_042137.1), ^*2*^*C. xerosis* (NR_026213.1), \**Bifidobacterium longum* (NR_145535.1), \**Corynebacterium diphtheriae* (NR_037079.1), ^1^*M. luteus* (NR_037113.1), ^*^*Cutibacterium acnes* (NR_145912.1), \**Streptomyces levis* (NR_041184.1), *Atopobiumminutum* (NR_044838.1), and \**Eggerthella lenta* (NR_037089.1) were used as Actinobacteria representatives. \**Anaplasma phagocytophilum* (NR_044762.1), \**Burkholderia cepacian* (NR_029209.1), ^1^*N. perflava* (NR_117694.1), \**Bilophila wadsworthia* (NR_176800.1), \**Legionella pneumophila* (NR_041742.1), ^1^*E. coli* (NR_024570.1), ^1^*Y. pseudotuberculosis* (NR_025158.1), ^*2*^*Klebsiella aerogenes* (NR_102493.2), ^*2*^*Pseudomonas putida* (NR_043424.1), ^*2*^*Stenotrophomonas maltophila* (NR_040804.1), ^*2*^*Vibrio natriegens* (NR_026124.1), and \**Vibrio cholerae* (NR_044050.1) were used as Proteobacteria representatives. \**Fusobacterium nucleatum* (NR_074412.1) was used as a Fusobacteria representative. \**Opitutus terrae* (NR_028890.1) and \**Akkermansia muciniphila* (NR_042817.1) were used as Verrucomicrobia representatives. MAFFT Version 7 (https://mafft.cbrc.jp/alignment/server/index.html) was used to align 16S rRNA sequences under a slow, iterative refinement method and to make the phylogenetic tree with 1000x Bootstrap. iTOL V. 6 (https://itol.embl.de/) was used for presentation of the phylogenetic tree.

### Antibodies, PAMPs, and Other Reagents

Antibodies against total STAT1 (#9172S), pS365-STING (mouse-reactive, #72971S), total STING (#13647), cGAS (#D3O8O), RIP2 (#4142), MAVS (#4983S), pS172-TBK1 (#D52C2), total TBK1 (rabbit background, #3504), total TBK1 (mouse background, #51872S), pS396-IRF3 (#4947S), total IRF3 (#4302S), and LRRC8A (#24979S) were purchased from Cell Signaling Technology. A second pS365-STING (mouse-reactive, #PA5-117271) was purchased from Thermo Fisher Scientific. The pY701-STAT1 antibody was purchased from BD (#612133). A mouse-reactive viperin antibody was purchased from EMD (#MABF106). The β-tubulin antibody was purchased from Developmental Studies Hybridoma Bank (#E7). The FLAG antibody was purchased from Sigma (#F1804-1MG). The *E. coli* antibody was purchased from Abcam (#137967).

*E. coli* LPS (serogroup O55:B5) was purchased from Enzo (ALX-581-013-L002). ADU-S100 (2′3′-c-di-AM(PS)2 (Rp, Rp), #tlrl-nacda2r), 2′3′-cGA (#tlrl-nacga23-1), 3′3′-cGG (#tlrl-nacdg), 3′3′-cAA (#tlrl-nacda), lipoteichoic acid (#tlrl-pslta), Poly(I:C) (#tlrl-pic), 5′ppp-dsRNA (#tlrl-3prna), N-glycolylated muramyl dipeptide (#tlrl-gmdp), DMXAA (#tlrl-dmx), and R848 (#tlrl-r848-5) were purchased from Invivogen. Sendai virus (SeV, Cantell Strain) supplied as allantoic fluid was sourced from Charles River Laboratory (#10100774). Calf thymus DNA (CT DNA) was purchased from Sigma (#D1501-100MG).

Recombinant IFNβ was purchased from BioLegend (#581302). Cytochalasin D (#C8273-5MG) was purchased from Sigma. H-151 (#inh-h151) and Bafilomycin A1 (#tlrl-baf1) were purchased from Invivogen. DCPIB was purchased from Tocris (#1540). LDH Cytotoxicity Assay kit (#C20301) was purchased from Thermo Fisher Scientific.

### Cell Culture and Primary Macrophage Differentiation

Immortalized bone marrow macrophages (iBMDMs), M-CSF-secreting L929 fibroblasts, and HEK293T cells were cultured at 37°C with 5% CO_2_ in DMEM (Gibco #11995-065) supplemented with 10% fetal bovine serum (FBS, Gibco #26140-079), 1x penicillin/streptomycin antibiotics (Gibco #15140-122), 1x L-glutamine (#25030-081), and 1x sodium pyruvate (#11360-070), herein referred to as “complete DMEM” (cDMEM). For passaging, iBMDMs were lifted with 1xPBS (Gibco#10010-023) supplemented with 2.5 mM EDTA (Invitrogen #15575-038) and replated with cDMEM at a dilution ratio of 1:10 on TC-treated plastic. WT and *Myd88*^*-/-*^*Trif*^*-/-*^ iBMDMs were previously generated in our lab (*80*). For passaging L929 or HEK293T cells, the cells were lifted with Trypsin-EDTA (Thermo Fisher Scientific #25200-114), trypsin inactivated with an equal volume of cDMEM, 10 percent of cells centrifuged at 400xg 5 min, and the cell pellet resuspended in cDMEM and plated on TC-treated plastic.

For primary bone marrow-derived macrophages, L929 cells were grown to confluency, incubated for 48 hrs, and M-CSF-containing supernatants were then collected and passaged through a 0.22 μm filter. The femurs and tibiae of dead male or female C57BL/6 mice (6-10 weeks old, Jackson Laboratories) were cleaned, cut with scissors, and centrifuged at 800xg for 5 min to pellet the bone marrow. The bone marrow pellet was then resuspended in 1xPBS supplemented with 1x antibiotic/antimycotic (Thermo Fisher Scientific #15240112), passaged through a 70 μm filter, and centrifuged at 400xg for 5 min. The bone marrow pellet was resuspended in differentiation medium (*i*.*e*., DMEM supplemented with 10% FBS, 1x antibiotic/antimycotic, and 30% L929 supernatant) at a concentration of 0.4×10^6^ cells/mL, plated on non-TC-treated 10cm dishes (10mL per plate), and incubated at 37°C with 5% CO_2_ (Day 0). At day 3, 5mL of differentiation medium was added. At day 6, cells were lifted with 1xPBS supplemented with 2.5 mM EDTA, centrifuged at 400xg 5 min, and the pellet resuspended in differentiation medium with reduced L929 supernatant (*i*.*e*., DMEM supplemented with 10% FBS, 1x antibiotic/antimycotic, and 5% L929 supernatant) at a concentration of 0.5×10^6^ cells/mL. 1mL BMDMs were plated in each well of a 24-well TC-treated plate and allowed to adhere overnight at 37°C with 5% CO_2_. Prior to infection, BMDM supernatants were aspirated from each well and 1mL DMEM supplemented with 10% FBS and 5% L929 supernatant was added to each well and remained present throughout the experiment.

For experiments using defined hypotonic medium, the hypotonic solution was made with endotoxin-free molecular grade water (Thermo Fisher Scientific #10-977-023) and the following defined solutes: 16.17 mM NaCl (Sigma #S8776), 6 mM KCl (Sigma #P9327-100ML), 1 mM MgCl_2_ (Thermo Fisher Scientific #AM9530G), 1.5 mM CaCl_2_ (Sigma, #C-34006), 10 mM glucose (Thermo Fisher Scientific #A2494001), 10 mM Hepes pH7.4 (Thermo Scientific Fisher #J16924.AP). For all other experiments, hypotonic conditions were created by diluting DMEM (Gibco #11995-065) 1:2 or 1:4 with endotoxin-free molecular grade water (Thermo Fisher Scientific #10-977-023).

### Bacterial Strains and General Growth Conditions

Strains used in this study are summarized in **Supplemental Table 1**. WT strains of *L. innocua* (#33091), *L. lactis* (#19435), *L. plantarum* (#14917), *M. luteus* (#4698), *E. faecalis* (#10100), *B. subtilis* (#6633), *S. capitis* (#146), *S. caprae* (#55133), *S. lugdunensis* (#49576), *S. saprophyticus* (#15305), *L. acidophilus* (#4356), *L. delbrueckii* (#11842), *E. casseliflavus* (#700327), *E. saccharolyticus* (#43076), *S. salivarius* (#13419), *S. mutans* (#25175), *S. gallolyticus* (#9809), *P. damnosus* (#11842), *C. pseudodiphtheriticum* (#10700), *C. xerosis* (#373), *K. aerogenes* (#13048), *P. putida* (#12633), *S. maltophila* (#13637), *V. natriegens* (#14048), S. carnosus (#51365), and *N. perflava* (#14799) were purchased from ATCC. WT *S. aureus* strain SA113 (ATCC #35556) was a gift from David Underhill. WT parent USA300, *ΔgdpP, ΔdacA S. aureus* were a gift from Angelika Gründling (*31, 32*). WT parent USA300 and *Δtoxins* were a gift from Victor Torres (*54*). WT *S. epidermidis* was a gift from Paula Watnick (*81*). WT strains of *B. fragilis* and *B. thetaiotaomicron* were a gift from Dennis Kasper (*82*) and Andy Goodman (*83*), respectively. These *Bacteroides* possess modified LPS that poorly stimulates TLR4 (*84, 85*). *L. rhamnosus* (strain LMS2-1), *L. reuteri* (strain CF48-3A), *E. coli* strain Nissle, *L. gasseri* (strain MV-22), *L. murinus* (strain WG mouse), *L. johnsonii* (strain CB mouse) were gifts from Dennis Kasper. WT K12 *E. coli* is strain MG1655 and used as the primary model *E. coli* in this study. *Y. pseudotuberculosis* used in this study was a gift from Igor Brodsky and is a plasmid-cured IP2666 strain that carries the mCD1 plasmid (*55*); mCD1 is modified to carry all the genes necessary to build a type 3 secretion system but lacks all effector and chaperone genes (*56*). Rosetta(DE3)pLysS and ClearColi competent *E. coli* (*i*.*e*., *E. coli* expressing Lipid IVa-containing LPS that is less able to stimulate murine TLR4 (*58*)) were purchased from Novagen (#70956) and Biosearch Technologies (#60810-1), respectively, and used as the parent strains of engineered *E. coli*. WT L. monocytogenes (strain 10403S) and *ΔhlyΔplcAΔplcB* (parent strain 10403S, designated as DP-L2319) were a gift from Darren Higgins.

*S. aureus* strain SA113, *S. capitis, S. caprae, S. epidermidis*, and *E. coli* (K12, ClearColi, Rosetta, and Nissle) were cultured on Luria Broth (LB, Invitrogen #12-780-052; Miller LB for ClearColi, Sigma #L3152-1kg) agar (1.5% w/v, BD #214010) at 37°C. *L. innocua, L. monocytogenes* (WT and *ΔhlyΔplcAΔplcB*), *C. pseudodiphtheriticum, C. xerosis, S. salivarius, S. mutans, S. gallolyticus, E. casseliflavus, E. saccharolyticus*, and *E. faecalis* were cultured on Brain Heart Infusion (BHI, BD #237500) agar at 37°C. *B. subtilis* was cultured on BHI agar at 30°C. *L. lactis* was cultured on BHI agar at 37°C with 5% CO_2_. *L. plantarum, L. rhamnosus, L. reuteri, L. gasseri, L. murinus, L. johnsonii, L. acidophilus*, and *L. delbrueckii* were cultured on lactobacilli MRS broth (BD #288130) agar at 37°C with 5% CO_2_. *M. luteus*, USA300 background *S. aureus, S. carnosus, S. lugdunensis*, and *S. saprophyticus* were cultured on tryptic soy broth (TSB, BD #211825) agar at 30°C (for 48 hrs for *M. luteus*) or 37°C (for Staphylococci). *For ΔdacA S. aureus*, TSB (BD #211825 with a total of 0.0855M NaCl) was supplemented with NaCl to 0.8M final concentration. *Y. pseudotuberculosis* (parental bacteria and *YP::dacA+*) was cultured on 2xYT broth (0.5% w/v NaCl, 1.6% w/v tryptone, 1% w/v yeast extract) agar at 26°C. *N. perflava* was cultured on Gonococcal medium base (GCB, BD #DF0289-17-3) plus Kellogg’s supplements (*86*) at 37°C with 5% CO_2_.

*S. maltophila, K. aerogenes*, and *P. putida* were cultured on Nutrient Broth (BD #234000) agar at 37°C, 30°C, or 26°C, respectively. *V. natriegens* was cultured on Nutrient Broth agar supplemented with 1.5% NaCl at 26°C. *P. damnosus* was cultured on MRS agar at 26°C. *B. fragilis* and *B. thetaiotaomicron* were cultured on trypticase soy-agar plates with 5% v/v sheep blood (Thermo Fisher Scientific #B21239X) at 37°C while contained within an anaerobic chamber (BD GasPak #260683), as per manufacturer’s instructions.

### CRISPR-mediated Knockout Generation

To generate gene knockouts in WT and *Myd88*^*-/-*^*Trif*^*-/-*^ iBMDMs via CRISPR-Cas9, guide oligos targeting genes of interest were designed with CRISPick (https://portals.broadinstitute.org/gppx/crispick/public) and are listed in **Supplemental Table 2**. One guide was previously published (*87*). Guide sequences were incorporated into sense and antisense oligos with the appropriate Esp3I overhangs on the 5′ end (*e*.*g*., sense oligo: 5′ CACC<u>G</u>Nx20 3′, where CACC corresponds to the 5’ Esp3I overhang, <u>G</u> helps with guide transcription, and Nx20 is the 20 nucleotide guide sequence) and 3′ end (*e*.*g*., antisense oligo: 5′ AAACNx20<u>C</u> 3′, where AAAC is the 3’ Esp3I overhang, Nx20 is the reverse complement of the 20 nucleotide guide sequence, and <u>C</u> base pairs with G in the sense oligo). To phosphorylate and anneal guide oligos, 2μL each of 50 μM sense and 50 μM antisense oligos were mixed with 1X T4 ligase buffer (NEB #B0202S) and 0.5μL T4 polynucleotide kinase (NEB #M0201S) for a total 10 μL reaction. The reaction mixture was incubated in a thermal cycler with the following steps: 1) 37°C for 30 min, 2) 95°C for 5 min, 3) 2°C/sec decrease to 85°C, 4) 85°C for 1 min, and 5) 0.1°C/sec decrease to 25°C. Phosphorylated, annealed oligos were diluted 1:20 in molecular grade H_2_O and then ligated into Esp3I-digested pLentiCRISPR V2.0 (Addgene #52961). Ligation reactions were then transformed into Top10 competent *E. coli* (Thermo Fisher Scientific #C404010) and selected with 100 μg/mL ampicillin.

To transduce iBMDMs with lentivirus, 2.5×10^6^ HEK293T cells were seeded into a 10 cm TC-treated dish and incubated overnight at 37°C with 5% CO_2_. HEK293T cells were then transfected with a mixture of 10 μg of pLentiCRISPR containing the CRISPR guide of interest, 10 μg of psPAX2 lentiviral packing plasmid, and 2.5 μg pCMV-VSV-G with Opti-MEM (Thermo Fisher Scientific # 31985062) and Lipofectamine 2000 (Thermo Fisher Scientific #11668019) according to manufacturer’s instructions. After an overnight incubation of the transfection at 37°C with 5% CO_2_, supernatants were aspirated from transfected cells, 6 mL cDMEM added, and cells incubated for 24 hrs at 37°C with 5% CO_2_. 0.5×10^6^ iBMDMs were plated per well of a 6-well TC plate and incubated overnight at 37°C with 5% CO_2_. 24 hr viral particle-laden supernatants were harvested, centrifuged at 400xg 5 min, passaged through a 0.45 μm filter, and mixed with 0.05% v/v Polybrene (Millipore TR-1003-G). iBMDM supernatants were aspirated and cells exposed to the prepared virus mixture and centrifuged at 1250xg 1 hr at 30°C. Transduced cells were allowed to rest with added cDMEM for 24 hrs, and then cells were subjected to one more round of viral transduction. After

1. hrs rest, cells were exposed to cDMEM supplemented with 10 μg/mL puromycin (Gibco #A11138-03) for one week to select for transduced cells. To harvest single CRISPR knockout clones, puromycin-selected cells were diluted to 1 cell per well in a 96-well TC-treated plate and allowed to propagate. Where possible, clones were confirmed as knockouts via loss of protein expression by immunoblot. In the case of IFNAR1 knockouts, clones were screened by the absence of IP-10 secretion in response to recombinant IFNβ. In contexts where antibodies do not exist, clones were confirmed by genomic DNA sequencing and analyzed for knockout status by ICE (https://ice.synthego.com/).

### Cloning

To create engineered *E. coli*, a gBlock of DacA (WP_000347896.1) was designed with a 5′ BamHI site, a C-terminal 3x-FLAG tag, and a 3′ EcoRI site, and was codon optimized for expression in *E. coli*. The gBlock was digested with BamHI and EcoRI and ligated into pET28a(+), which contains an IPTG-inducible lac promoter. Transformants were selected with 50 μg/mL kanamycin. Rosetta(DE3)pLysS or Clearcoli *E. coli* were then transformed with a modified pET28a(+) vector lacking a His-Tag (*88*), and transformants selected with 50 μg/mL kanamycin and 25 μg/mL chloramphenicol. To create YP::*dacA+, dacA-1xFLAG* was PCR amplified from pET28a(+)-*dacA-3xFLAG* with the forward primer 5′ ACCGC<u>CATATG</u>GATTTCTCTAACTTCTTCC 3′ (NdeI site is underlined) and the reverse primer 5′ CAGCC<u>GGATCC</u>CTACTTATCGTCGTCATC 3′ (BamHI site is underlined) and subcloned into the pET16 with NdeI and BamHI restriction sites. *Y. pseudotuberculosis* was electroporated with the resulting pET16-*dacA-1xFLAG* as described in (*89*) and clones selected for with 100 μg/mL ampicillin. To mutate CDN synthase codons, Q5 mutagenesis (NEB # E0554S) on pDNA was performed according to the manufacturer’s instructions using primers designed via NEBaseChanger (https://nebasechanger.neb.com/).

For macrophage gene reconstitution experiments, gBlocks were designed with a 5′ XhoI site, a C-terminal 3x-FLAG tag or left untagged as in the case for *Sting* constructs, and a 3′ NotI site, and gBlocks were codon optimized for expression in *M. musculus*. Refer to **Supplemental Table 2** for gene sequence IDs. gBlocks were digested with XhoI and NotI and ligated into the retrovirus vector pMSCV-IRES-GFP, which is used for stable reconstitution of macrophages and GFP signal as a proxy for expression of the gene of interest. Transformants were selected for with 100 μg/mL ampicillin. For doxycycline-inducible gene expression, constructs were PCR amplified from pMSCV-IRES-GFP pDNA. For *Lrrc8a-3xFLAG*, the forward primer 5′ ATCCATCGAT<u>ACGCGT</u>GCCACCATGATACCCG 3′ (MluI site is underlined) and the reverse primer 5′ GCCCGCCGGC<u>GCGGCCGC</u>CTATTTATCGTCGTC 3′ (NotI site is underlined) were used. For *Slc46a2-3xFLAG*, the forward primer 5′ TCTTATACTT<u>GGATCC</u>GCCACCATGGGTCCCGGT GG 3′ (BamHI site is underlined) and the reverse primer 5′ GCGCGGCCGC<u>ACGCGT</u>CTACTTATCGTCGTC 3′ (MluI site is underlined) were used. Digested PCR amplicons were then ligated into pRetro-TRE3g (Takara). To mutate the codons of the *Sting* gene, Q5 mutagenesis of pDNA was performed as above.

Retroviral transduction of iBMDMs was performed as done with lentiviral transduction above with the exception that 10 μg of pMSCV-IRES-GFP constructs, pRetroX-Tet3G (Takara), or pRetro-TRE3g constructs was mixed with 3 μg pCMV-VSV-G and 6 μg pCL-ECO retroviral packing plasmid for HEK293T transfections. For pMSCV-IRES-GFP, iBMDMs with expression of the gene of interest were enriched by FACS by sorting for GFP positive cells. For doxycycline-inducible cells, iBMDMs were first transduced with pRetroX-Tet3G for reverse tetracycline controlled transactivator expression, and transduced iBMDMs were enriched by selection with 1.5 mg/mL G-418. Then, pRetroX-Tet3G-containing iBMDMs were then transduced with pRetro-TRE3g constructs and transduced cells enriched by selection with 1.5 mg/mL G-418 and 10 μg/mL puromycin. 0.5 μg/mL doxycycline was used for expression of the gene of interest in pRetroX-Tet3G/pRetro-TRE3g-transduced cells.

### Bacterial Infections and Treatments with Stimuli or Inhibitors

To prepare bacteria for infection, *S. aureus* (SA113), *S. capitis, S. caprae, S. epidermidis*, and *E. coli* (K12, ClearColi, Rosetta, and Nissle) colonies were inoculated into liquid LB (Miller LB for ClearColi) and grown overnight aerobically at 37°C. USA300 strains of *S. aureus, S. carnosus, S. lugdunensis*, and *S. saprophyticus* colonies were inoculated into liquid TSB and grown overnight aerobically at 37°C with the exception that *ΔdacA* was grown in liquid TSB with a total of 0.8M NaCl. For Rosetta *E. coli*, 25 μg/mL chloramphenicol (to select for pLysS) and/or 50 μg/mL kanamycin (to select for pET28a(+) constructs) were used throughout growth. For ClearColi, 50 μg/mL kanamycin (to select for pET28a(+) constructs) were used throughout growth. To induce protein expression, *E. coli* were treated with isopropyl β-D-1-thiogalactopyranoside (IPTG, Thermo Fisher Scientific # 15529019) at indicated concentrations (.01-1 mM) for 3 hrs just prior to infection. *L. innocua, L. monocytogenes, C. pseudodiphtheriticum, C. xerosis, S. salivarius, S. mutans, S. gallolyticus, E. casseliflavus, E. saccharolyticus*, and *E. faecalis* colonies were inoculated into liquid BHI and grown overnight aerobically at 37°C. *L. lactis* colonies were inoculated into liquid BHI supplemented with 0.042% w/v NaHCO_3_, and grown overnight aerobically at 37°C.

*L. plantarum, L. rhamnosus, L. reuteri, L. gasseri, L. murinus, L. johnsonii, L. acidophilus*, and *L. delbrueckii* colonies were inoculated into liquid MRS supplemented with 0.042% w/v NaHCO_3_, and grown overnight aerobically at 37°C. *B. subtilis* and *M. luteus* colonies were inoculated into liquid BHI and TSB, respectively, and grown overnight aerobically at 30°C. Approximately 15 *N. perflava* colonies were inoculated onto GCB agar, grown overnight at 37°C with 5% CO_2_, and stationary phase bacteria swabbed into liquid gonococcal media (1.5% w/v proteose peptone #3, 0.4% w/v K_2_HPO_4_, 0.1% w/v KH_2_PO_4_, and 0.1% w/v NaCl, pH 7.2) modified with Kellogg’s supplements and 0.042% w/v NaHCO_3_. *Y. pseudotuberculosis* colonies were inoculated into liquid 2xYT and grown overnight aerobically at 26°C. YP::*dacA+* bacteria were treated with 100 μg/mL ampicillin throughout growth. *S. maltophila, K. aerogenes*, and *P. putida* colonies were inoculated into liquid Nutrient Broth and grown overnight aerobically at 37°C, 30°C, or 26°C, respectively. *V. natriegens* colonies were inoculated into liquid Nutrient Broth (1.5% NaCl supplement) and grown overnight aerobically at 26°C. *P. damnosus* colonies were inoculated into liquid MRS and grown overnight at 26°C.

Stationary phase bacteria in liquid cultures (*i*.*e*., overnight liquid cultures or agar cultures from above) were then subjected to two back dilutions (for *M. luteus* and *P. damnosus*, one back dilution) to enrich for highly viable, logarithmic phase cultures that were used for infections. For *Y. pseudotuberculosis*, overnight cultures were subsequently back diluted and induced for expression of the type 3 secretion system as previously described (*90*). For YP::*dacA+* experiments, parent and YP::*dacA+* were grown as before except in LB. For non-inducing type 3 secretion system conditions, parent and YP::*dacA+* overnight cultures were back diluted into LB and grown for 3 hrs aerobically at 37°C; all conditions received 1 mM IPTG during this 3 hr period. Approximately 15 *B. fragilis* and *B. thetaiotaomicron* colonies were inoculated onto agar as described above for 48 hrs anaerobically prior to being swabbed into BHI liquid media for its immediate use in infections.

At the time of infection, bacteria were centrifuged at 10,000xg 3 min, supernatant discarded, and pellets suspended in DMEM supplemented with 10% heat-inactivated FBS, 1x L-glutamine (#25030-081), 1x sodium pyruvate (#11360-070), herein referred to as “infection DMEM” (iDMEM). Bacteria suspended in iDMEM were plated to confirm multiplicity of infection (MOI). To prepare iBMDMs for infections, cultured iBMDMs were centrifuged at 400xg 5 min, the pellet resuspended to a concentration of 0.5×10^6^ cells/mL in iDMEM, and 1 mL or 0.2 mL cells was seeded into each well of a 24-well or 96-well TC-treated plate, respectively. Seeded iBMDMs were allowed to adhere for at least 4 hrs at 37°C with 5% CO_2_. To initiate a synchronous infection, adherent iBMDMs were placed on ice for 5 min, supernatants aspirated, and then cells in the 24-well plate setup were exposed to 1 mL bacteria at the indicated MOI or 1 mL PAMPs at the indicated concentration. Cells were centrifuged at 400xg 5 min 4°C to initiate bacterial contact with iBMDMs. The infection was incubated for 1 hr at 37°C with 5% CO_2_ to allow for phagocytosis, after which infected supernatants were aspirated to remove non-adherent, extracellular bacteria. cDMEM containing 1x pen/strep was then added to eliminate remaining extracellular bacteria, and the infection was allowed to proceed at 37°C with 5% CO_2_ for the total indicated time. For experiments with inhibitors or BMDMs, pen/strep at a final concentration of 1x was spiked into cells to inactivated extracellular bacteria.

In cases where bacteria were inactivated prior to infection, pellets containing 1×10^8^ CFU (as grown above) were resuspended in 1 mL 4% paraformaldehyde (PFA) in 1x PBS and incubated for 15 min at room temperature. PFA-inactivated bacteria were then centrifuged at 10,000xg 3 min, PFA supernatants removed, and pellets washed with 1 mL 1xPBS. After one more centrifugation step, the pellet of PFA-inactivated bacteria was resuspended in iDMEM and plated to confirm bacterial inactivation.

To determine recoverable CFU during infection, iBMDMs were infected with an MOI of 50 as above. The inoculum was serially diluted and plated for viable CFU to determine starting CFU/mL. After 1 h infection, supernatants were removed and replaced with cDMEM and pen/strep. After 4 hrs or 24 hrs, supernatants were removed from infected iBMDMs, and cells were lysed with 200 μL 1% saponin (Sigma #47036) per well of a 24-well plate for 10 min at 37°C. 800 μL LB were then added to each well and mixed vigorously. Lysates were serially diluted and plated for viable CFU.

For cell association experiments by flow cytometry, bacteria were labelled with 5-(and-6)-carboxyfluorescein diacetate, succinimidyl ester (CFSE) as in (*91*) and exposed to macrophages as above. After 2 hrs total incubation, macrophage supernatants were aspirated, cells washed once with 1 mL 1xPBS, and then cells lifted with 1xPBS with 2.5 mM EDTA. Cells were then centrifuged at 400xg 5 min, supernatants aspirated, and cells fixed with 1% PFA in 1xPBS. Data were acquired on a LSRFortessa™ cell analyzer (BD) and analyzed using FlowJo.

For experiments using bacterial conditioned supernatants, *S. aureus* and *E. coli::dacA+* were grown as above, except with the following modification at the final back dilution step: 2 mL mid-log phase culture was centrifuged at 10,000xg 3 min, the pellet resuspended in 16 mL iDMEM (without phenol red) with 0.01 mM IPTG, and cultured aerobically for 3 hrs 37°C. Live bacteria were prepared as normally done above from these cultures. The remaining cultures were then equilibrated to an equivalent OD_600_ with iDMEM, centrifuged at 4,122xg 5 min, the supernatants passaged through a 0.22 μm filter, and 1 mL supernatants were applied to 0.5×10^6^ macrophages per well.

For SeV infection, SeV was diluted in DMEM supplemented with 1% FBS in a total volume of 100 μL per well (24-well plate format) for an MOI of 0.00001. iBMDMs were exposed to SeV for 1 hr at 37°C with 5% CO_2_ during which plates were gently rocked every 10 min. After 1 hr, supernatants were removed, replaced with 1 mL cDMEM, and infection incubated at 37°C with 5% CO_2_ for total indicated time.

All PAMPs were diluted in iDMEM to the indicated concentration before exposure to macrophages. For cytosolic transfections of PAMPs, 1 μg CT DNA, 100 μg CDNs (where indicated), or 0.5 μg 5′ppp-dsRNA was incubated with Opti-MEM and 3 μL Lipofectamine-2000 (Thermo Fisher Scientific #11668019) for a total of 100 μL for at least 20 min prior to infection. Supernatants from macrophages were aspirated and 900 μL iDMEM added to each well, followed by 100 μL drop-wise addition of PAMPs for a final concentration of 1 μg/mL CT DNA, 100 μg/mL CDNs, or 0.5 μg/mL 5′ppp-dsRNA per well.

For cytochalasin D treatment, iBMDMs were treated with vehicle (DMSO) or 10 μg/mL cytoD for 10 min prior to and throughout infection. For the H-151 STING inhibitor, macrophages were treated with vehicle (DMSO) or 0.5 μg/mL (iBMDMs) or 1 μg/mL (BMDMs) H-151 for 1 hr prior to and throughout infection. For LRRC8A inhibition via DCPIB, iBMDMs were treated with vehicle (DMSO) or 20 μM DCPIB for 30 min prior to and throughout infection. For phagosome acidification inhibition, iBMDMs were treated with vehicle (DMSO) or 100 μM Bafilomycin A1 for 30 min prior to and throughout infection.

For assessment of phagocytosis by immunofluorescence, *E. coli*-infected cells (MOI of 10) were stained as in (*92*). Briefly, 10 min synchronously infected cells were fixed with 4% PFA for 15 min, washed 3 times with 1xPBS for 5 min each, blocked with 2% normal goat serum-1xPBS Jackson Laboratory (#005-000-121) for 1 hr, stained with anti-*E. coli* antibody (1:50 dilution) in 2% normal goat serum-1xPBS overnight at 4°C to label extracellular bacteria, washed as before, stained with goat anti-rabbit Alexa Flour 568 (Thermo Fisher Scientific #A11011) for 1 hr, and washed as before. Cells were then fixed again, washed, and permeabilized with 2% normal goat serum-1xPBS with 0.2% saponin for 1 hr. Cells were then stained with anti-*E. coli* antibody (1:50 dilution) in 2% normal goat serum-1xPBS with 0.2% saponin overnight at 4°C to label total bacteria, washed as before, stained with goat anti-rabbit Alexa Flour 488 (Thermo Fisher Scientific #A11008) and Alexa Flour phalloidin-647 (Thermo Fisher Scientific #A22287) for 1 hr, washed as before, exposed to DAPI (Biotium #40043) for 5 min, rinsed with water, and mounted with prolong antifade (Thermo Fisher Scientific #P36980). Images were acquired using a Zeiss 880 laser scanning confocal microscope.

### ELISA, immunoblotting, and quantitative reverse transcription polymerase chain reaction (qRT-PCR)

Macrophages exposed to bacteria and PAMPs were centrifuged at 400xg 5 min prior to analyses. Conditioned supernatants were collected, stored at - 20°C, and assayed by ELISA using IFNβ (Invivogen #luex-mifnbv2), IP-10 (R&D #DY466), and TNFα (Invitrogen #88-7324-88) kits, according to the manufacturer’s instructions. ELISA absorbance and luminescent readings were measured with a Tecan Spark 10M Multi-Mode Plate Reader. For 3′3′-cAA ELISA (Cayman Chemical #501960), bacteria were grown as if to be used for infections and 10×10^8^ CFU were pelleted as above. The bacterial pellets were lysed with 200 μL bacterial protein extraction reagent (Thermo Fisher Scientific #78243), and lysates were passaged through a 22g needle four times to reduce viscosity. Lysates were centrifuged at 10,000xg for 5 min and supernatants transferred to a fresh tube to remove any cell debris. Lysates were then applied to the 3′3′-cAA ELISA, according to the manufacturer’s instructions.

Protein lysates were prepared by adding 250 μL 1x sample buffer (12 mM Tris-HCl pH 6.8, 0.4% w/v SDS, 5% v.v glycerol, 0.02% w/v bromophenol blue, 1% v/v β-mercaptoethanol) per well of macrophages. Protein lysates were then boiled for 5 min prior to separation via Tris-Glycine SDS-PAGE gels. Proteins were then transferred onto PVDF membranes, which were subsequently blocked with 5% BSA in 1xTBS with 0.1% Tween-20 (TBS-T). Antibodies were diluted in 1% BSA TBS-T, according the manufacturer’s recommendations (except, anti-pS365-STING from Cell Signaling Technology, which was diluted 1:100). Blots were incubated with primary antibodies overnight at 4°C, and blots were incubated with secondary antibodies for 1 hr at RT. Blots were washed 5x with TBS-T before and after the secondary antibody incubation. Membranes were incubated with SuperSignal West Pico or Femto Chemiluminescent substrate (Thermo Fisher Scientific), and images of blots were acquired on a ChemiDoc XRS+ System (BioRad).

For RNA, macrophages were lysed with 300 μL lysis buffer (from Thermo Fisher Scientific #12183025) with 1% v/v β-mercaptoethanol. Total RNA was isolated (Thermo Fisher Scientific kit #12183025) and treated with DNaseI (Thermo Fisher Scientific #EN0521), according to the manufacturer’s instructions. 30 ng of RNA was used per well (of a 384-well plate format) for detection. Reverse transcription and PCR was performed on a CFX384 Touch Real-Time PCR detection system (BioRad) using the Taqman RNA-to-CT 1-Step kit (Thermo Fisher Scientific #4392938), according to manufacturer’s instructions. Genes were detected by Taqman probes (Mm01277042_m1 for *Tbp*; Mm00439552_s1 for *Ifnb1*; Mm00491265_m1for *rsad2*; and Mm99999068_m1 for *Tnf*; Mm00446220_m1 for *Slc19a1*; Mm00546630_m1 for *Slc46a1*; Mm00498614_m1 for *Slc46a2*; Mm00503612_m1 for *Scl46a3*; Mm00624918_m1 for *Lrrc8a*; Mm01206934_m1 for *Lrrc8b*; Mm00505716_m1 for *Lrrc8c*; Mm07304576_m1 for *Lrrc8d*; Mm01719515_m1 for *Lrrc8e*).

### Data Representation and Statistics

All values, unless otherwise indicated, represent the mean ± the standard error of the mean (SEM) of at least 3 independent replicates. Heat map values represent the mean of at least *n* = 3 biological replicates. All shown immunoblots and micrographs are representative of at least *n* = 3 biological replicates. A two tailed student’s *t*-test was performed to determine significance, which was defined as a *p*-value less than 0.05.

## Acknowledgements

We would like to thank Igor Brodsky, David Underhill, Paula Watnick, Dennis Kasper, Andy Goodman, Angelika Gründling, Victor Torres, and Darren Higgins for bacteria used in this study. We would also like to thank all members for the Kagan Laboratory for their helpful discussions and support.

S.A.R. was supported by postdoctoral fellowship from the Jane Coffin Childs Memorial Fund for Medical Research. This work was supported by NIH grants AI133524 (J.C.K.), AI116550 (JCK), and P30DK34854 (JCK).

## Supplementary Information

**Supplemental Figure 1.**
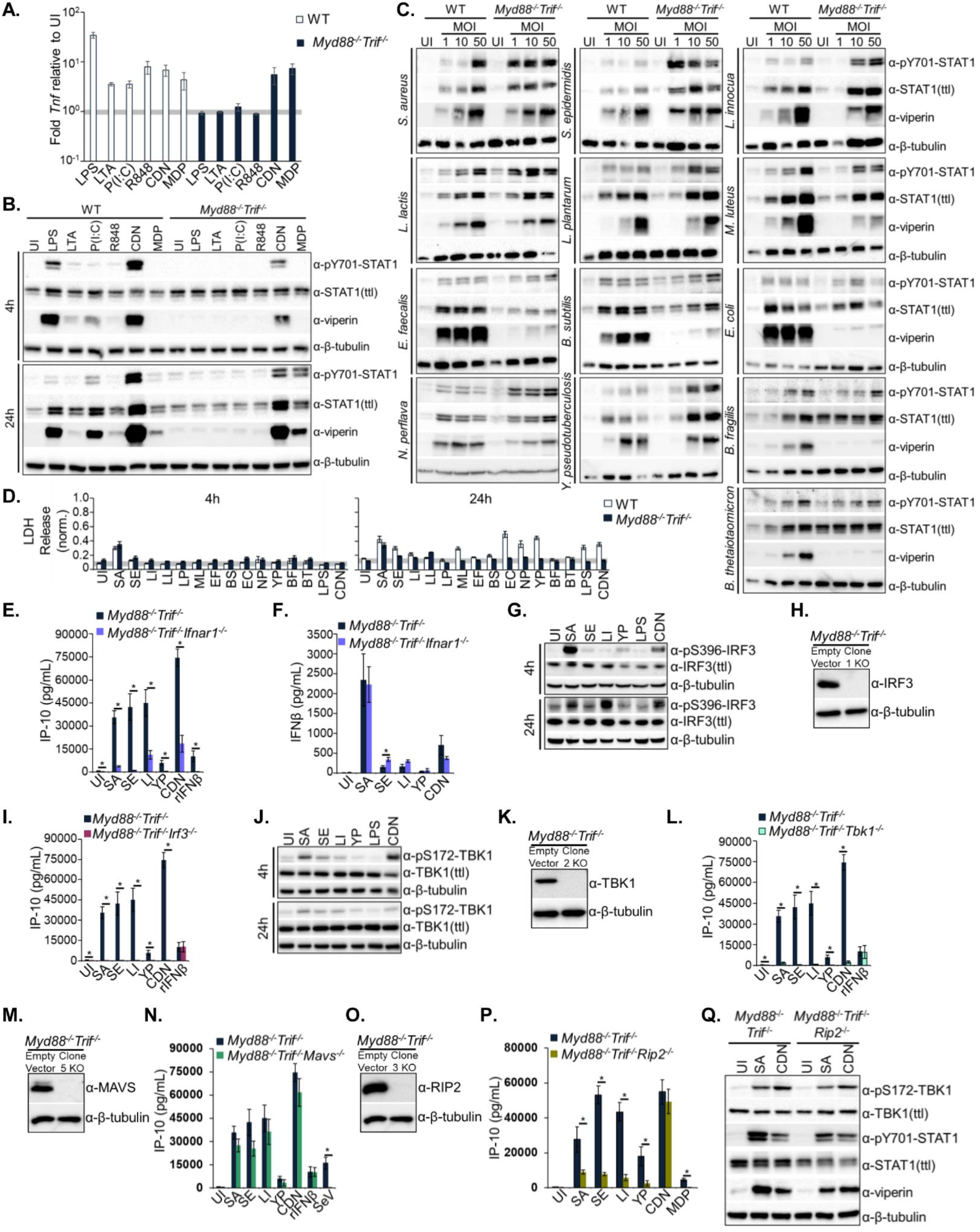
Immune responses in WT and *Myd88*^*-*/-^*Trif*^*-*/-^ macrophages to bacteria and PAMPs, related to Figure 1. **(A)** WT and *Myd88*^*-*/-^*Trif*^*-*/-^ iBMDMs were exposed to LPS (1 μg/mL; TLR4 agonist), lipoteichoic acid (“LTA”, 1 μg/mL; TLR2 agonist), poly(I:C) (“P(I:C)”, 50 μg/mL; TLR3 agonist), R848 (5 μg/mL; TLR7/8 agonist), ADU-S100 (“CDN”, 1 μg/mL; STING agonist), and N-glycolyl-muramyl dipeptide (“MDP”, 1 μg/mL, NOD2 agonist). 24 h RNA lysates were analyzed for fold *Tnf* transcriptional responses as in **Fig. 1C**. **(B)** WT and *Myd88*^*-*/-^*Trif*^*-*/-^ iBMDMs were treated as in **Suppl. Fig. 1A**. 4 h (top) and 24 h (bottom) protein lysates were analyzed by immunoblot for pSTAT1, total STAT1, viperin, and β-tubulin. **(C)** WT and *Myd88*^*-*/-^*Trif*^*-*/-^ iBMDMs were untreated (uninfected, “UI”) or exposed to bacteria at increasing MOIs. At 24 h post-exposure, protein lysates were analyzed by immunoblot for pSTAT1, total STAT1, viperin, and β-tubulin. **(D)** WT and *Myd88*^*-*/-^*Trif*^*-*/-^ iBMDMs were treated as in **Fig. 1B**. 4 h (left) and 24 h (right) conditioned supernatants were removed and assayed for the presence of LDH. LDH release in supernatants is normalized to LDH from total cell lysate controls. **(E)** *Myd88*^*-*/-^*Trif*^*-*/-^ and *Myd88*^*-*/-^*Trif*^*-*/-^*Ifnar1*^*-*/-^ iBMDMs were treated as in **Fig. 1B** with the additional treatment of recombinant IFNβ (“rIFNβ”, 50 ng/mL). 24 h conditioned supernatants were analyzed by ELISA for IP-10. **(F)** *Myd88*^*-*/-^*Trif*^*-*/-^ and *Myd88*^*-*/-^*Trif*^*-*/-^*Ifnar1*^*-*/-^ iBMDMs were treated as in **Fig. 1B**. 24 h conditioned supernatants were analyzed by ELISA for IFNβ. **(G)** *Myd88*^*-*/-^*Trif*^*-*/-^ iBMDMs were treated as in **Fig. 1B**. 4 h (top) and 24 h (bottom) protein lysates were analyzed by immunoblot for phosphorylated IRF3 (pS396), total IRF3, and β-tubulin. **(H)** Protein lysates were collected from *Myd88*^*-*/-^*Trif*^*-*/-^ and *Myd88*^*-*/-^*Trif*^*-*/-^*Irf3*^*-*/-^ iBMDMs and analyzed by immunoblot for IRF3 and β-tubulin. **(I)** *Myd88*^*-*/-^*Trif*^*-*/-^ and *Myd88*^*-*/-^*Trif*^*-*/-^*Irf3*^*-*/-^ iBMDMs were treated as in **Suppl. Fig. 1E**, and 24 h conditioned supernatants were analyzed by ELISA for IP-10. **(J)** *Myd88*^*-*/-^*Trif*^*-*/-^ iBMDMs were treated as in **Fig. 1B**. 4 h (top) and 24 h (bottom) protein lysates were analyzed by immunoblot for phosphorylated TBK1 (pS172), total TBK1, and β-tubulin. **(K)** Protein lysates were collected from *Myd88*^*-*/-^*Trif*^*-*/-^ and *Myd88*^*-*/-^*Trif*^*-*/-^*Tbk1*^*-*/-^ iBMDMs and analyzed by immunoblot for TBK1 and β-tubulin. **(L)** *Myd88*^*-*/-^*Trif*^*-*/-^ and *Myd88*^*-*/-^*Trif*^*-*/-^*Tbk1*^*-*/-^ iBMDMs were treated as in **Suppl. 1E**, and 24 h conditioned supernatants were analyzed by ELISA for IP-10. **(M)** Protein lysates were collected from *Myd88*^*-*/-^*Trif*^*-*/-^ and *Myd88*^*-*/-^*Trif*^*-*/-^*Mavs*^*-*/-^ iBMDMs and analyzed by immunoblot for MAVS and β-tubulin. **(N)** *Myd88*^*-*/-^*Trif*^*-*/-^ and *Myd88*^*-*/-^*Trif*^*-*/-^*Mavs*^*-*/-^ iBMDMs were treated as in **Suppl. Fig. 1E**, with the additional treatment of Sendai Virus (“SeV”, MOI 0.00001). 24 h conditioned supernatants were analyzed by ELISA for IP-10. **(O)** Protein lysates were collected from *Myd88*^*-*/-^*Trif*^*-*/-^ and *Myd88*^*-*/-^*Trif*^*-*/-^*Rip2*^*-*/-^ iBMDMs and analyzed by immunoblot for RIP2 and β-tubulin. **(P)** *Myd88*^*-*/-^*Trif*^*-*/-^ and *Myd88*^*-*/-^*Trif*^*-*/-^*Rip2*^*-*/-^ iBMDMs were treated as in **Fig. 1B and Suppl. Fig. 1A**. 24 h conditioned supernatants were analyzed by ELISA for IP-10. **(Q)** *Myd88*^*-*/-^*Trif*^*-*/-^ and *Myd88*^*-*/-^*Trif*^*-*/-^*Rip2*^*-*/-^ iBMDMs treated as in **Fig. 1B**. 4 h protein lysates were analyzed for analyzed by immunoblot for pTBK1, total TBK1, pSTAT1, total STAT1, viperin, and β-tubulin.

**Supplemental Figure 2.**
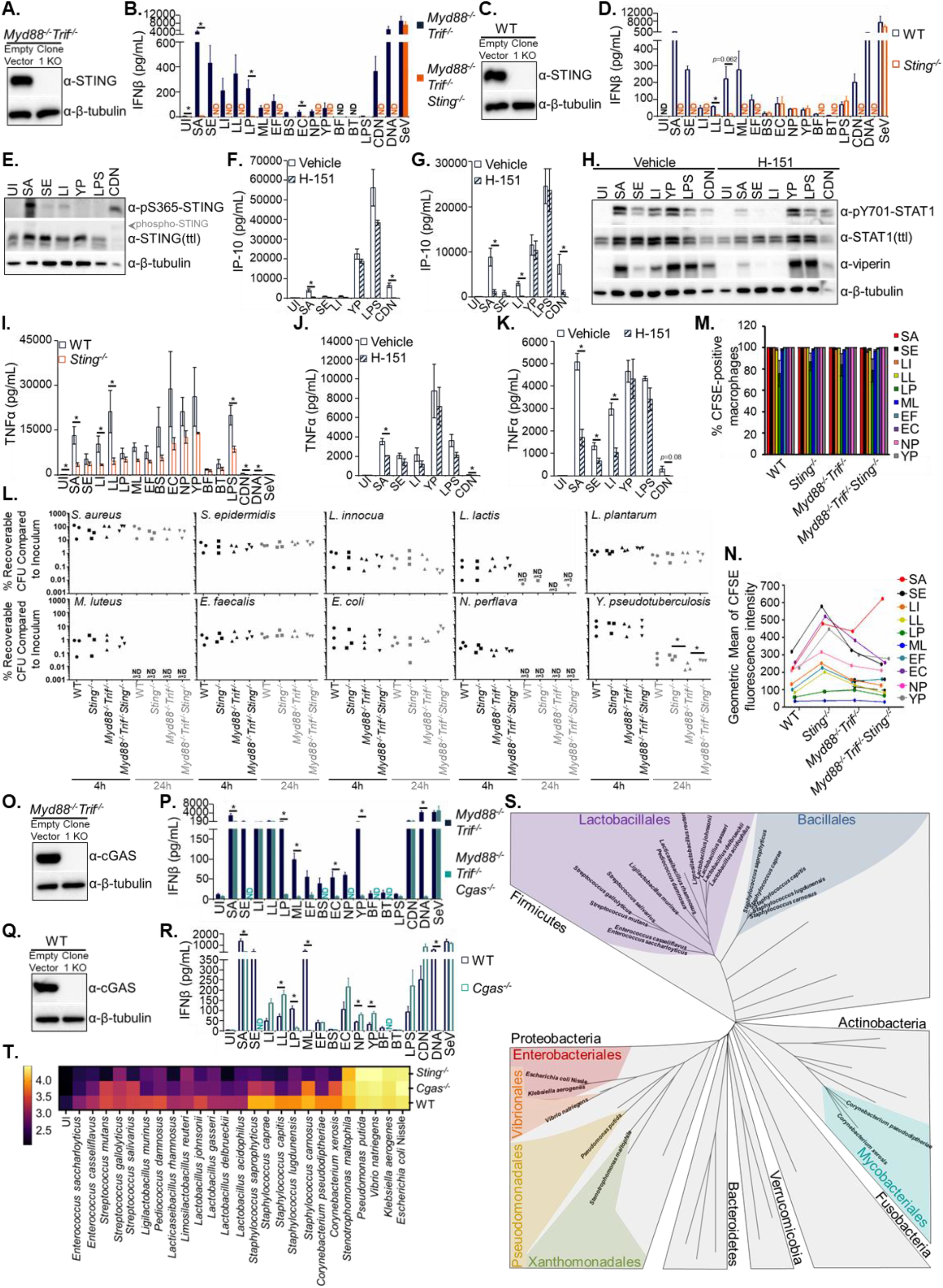
The role of STING in controlling TLR-independent IFN responses to bacteria in macrophages, related to Figure 2. **(A)** Protein lysates were collected from *Myd88*^*-*/-^*Trif*^*-*/-^ and *Myd88*^*-*/-^*Trif*^*-*/-^*Sting*^*-*/-^ iBMDMs and analyzed by immunoblot for STING and β-Tubulin. **(B)** 24 h conditioned supernatants from **Fig. 2B** were analyzed by ELISA for IFNβ. **(C)** Protein lysates were collected from WT and *Sting*^*-*/-^ iBMDMs and analyzed by immunoblot for STING and β-Tubulin. **(D)** 24 h conditioned supernatants from **Fig. 2C** were analyzed by ELISA for IFNβ. **(E)** WT BMDMs were treated as in **Fig. 1B**. 4 h protein lysates were analyzed by immunoblot for pSTING, total STING, and β-Tubulin. **(F)** WT iBMDMs were treated in the absence (vehicle) or presence of 0.5 μg/mL H-151 for 1 h prior to and throughout exposure to PAMPs and bacteria as in **Fig. 1B**. 4 h conditioned supernatants were analyzed by ELISA for IP-10. **(G)** WT BMDMs were treated in the absence (vehicle) or presence of 1 μg/mL H-151 for 1 h prior to and throughout exposure to PAMPs and bacteria as in **Fig. 1B**. 4 h conditioned supernatants were analyzed by ELISA for IP-10. **(H)** 4 h protein lysates from **Suppl. Fig. 2G** were analyzed by immunoblot for pSTAT1, total STAT1, Viperin, and β-Tubulin. **(I)** 24 h conditioned supernatants from **Fig. 2C** were analyzed by ELISA for TNFα. **(J)** 4 h conditioned supernatants from **Suppl. Fig. 2F** were analyzed by ELISA for TNFα. **(K)** 4 h conditioned supernatants from **Suppl. Fig. 2G** were analyzed by ELISA for TNFα. **(L)** WT (circles), *Sting*^*-*/-^ (squares), *Myd88*^*-*/-^*Trif*^*-*/-^ (triangles), and *Myd88*^*-*/-^*Trif*^*-*/-^*Sting*^*-*/-^ (upside-down triangles) iBMDMs were exposed to bacteria at an MOI of 50 and viable CFU assessed after 4 h (black color) and 24 h (grey color) (see methods). In some instances, CFU were not detected (as indicated in graphs, *n* = number of biological replicates below limit of detection). Percent recoverable CFU was calculated by dividing CFU at 4 h or 24 h by the CFU of the initial infection inoculum. For each bacterium at each time point, differences in values between genotypes were not statistically significant for any comparison, unless noted on the graph. **(M)** WT, *Sting*^*-*/-^, *Myd88*^*-*/-^*Trif*^*-*/-^, and *Myd88*^*-*/-^*Trif*^*-*/-^*Sting*^*-*/-^ iBMDMs were exposed to CFSE-labelled bacteria at an MOI of 50. After 2 h post-exposure, CFSE fluorescence was measured by flow cytometry. Shown is the percent (%) iBMDMs associated with CFSE-labelled bacteria. For each bacterium, differences in values between genotypes were not statistically significant for any comparison. **(N)** The geometric mean fluorescence intensity of CFSE-positive iBMDMs (*i*.*e*., infected cells) and a representative graph of 3 independent experiments from **Suppl. Fig. 2M**. For each bacterium, differences in values between genotypes were not statistically significant for any comparison. **(O)** Protein lysates were collected from *Myd88*^*-*/-^*Trif*^*-*/-^ and *Myd88*^*-*/-^*Trif*^*-*/-^*Cgas*^*-*/-^ iBMDMs and analyzed by immunoblot for cGAS and β-Tubulin. **(P)** 24 h conditioned supernatants from **Fig. 2D** were analyzed by ELISA for IFNβ. **(Q)** Protein lysates were collected from WT and *Cgas*^*-*/-^ iBMDMs and analyzed by immunoblot for cGAS and β-Tubulin. **(R)** 24 h conditioned supernatants from **Fig. 2E** were analyzed by ELISA for IFNβ. **(S)** Phylogenetic tree assembled as in **Fig. 1A**. Bacteria of the same color are within the same taxonomic order. Gram-negative bacteria are all from Proteobacteria. All other bacteria are gram-positives. **(T)** WT, *Cgas*^*-*/-^, and *Sting*^*-*/-^ iBMDMs were treated as in **Fig. 2B**. 24 h conditioned supernatants were analyzed by ELISA for IP-10. Data are expressed as in **Fig. 1B**.

**Supplemental Figure 3.**
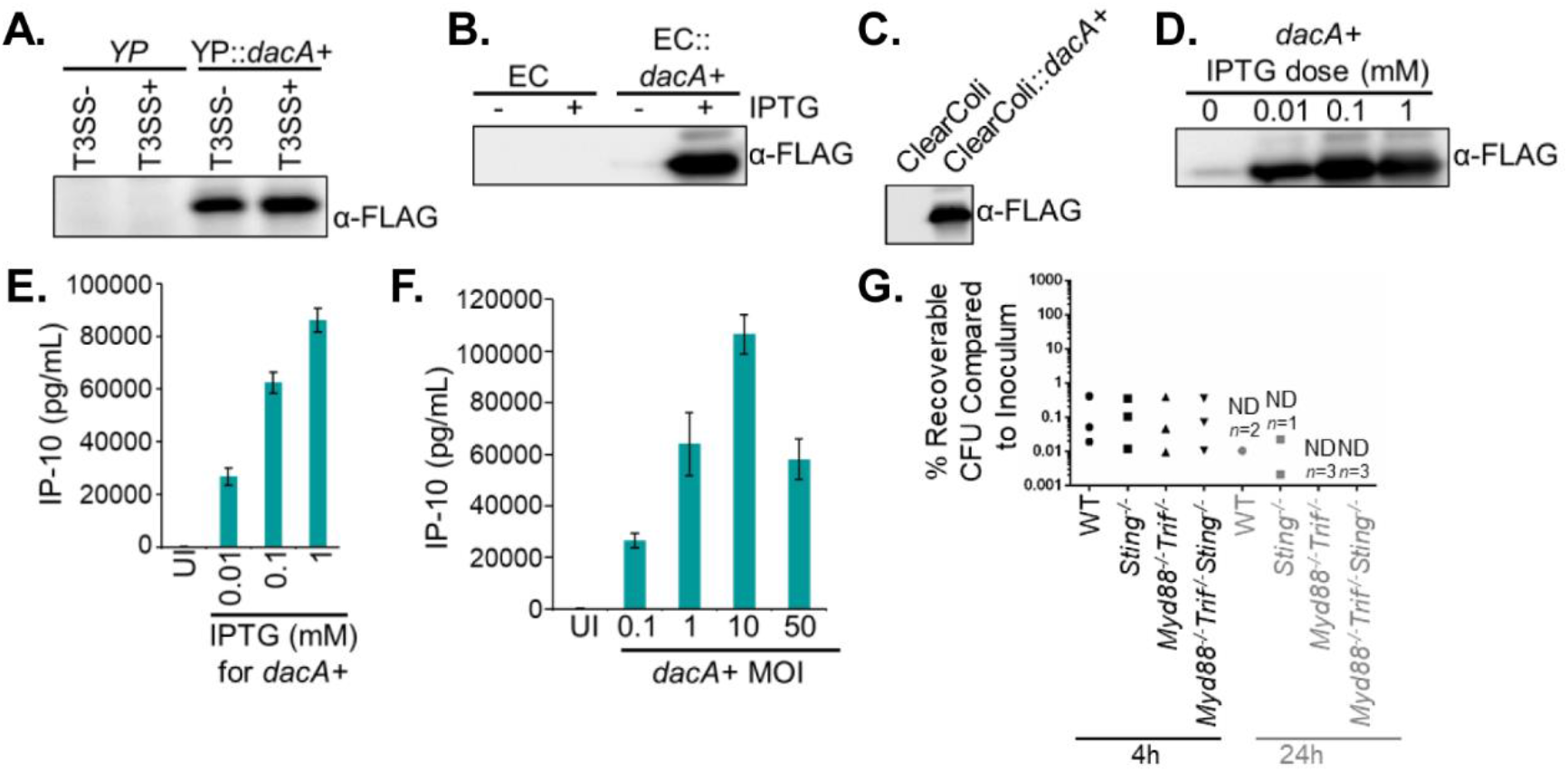
The role of virulence in STING activation to bacteria, related to Figure 3. **(A)** *Y. pseudotuberculosis* (“YP”) and *Y. pseudotuberculosis::dacA+* (“YP::*dacA+*”) were grown as in **Fig. 3D** to induce expression of DacA-1xFLAG. Protein lysates from bacteria were analyzed by immunoblot for FLAG protein. **(B)** *E. coli* and *E. coli::dacA+* were untreated (“-”) or treated (“+”) with 1 mM IPTG to induce expression of DacA-3xFLAG. Protein lysates from bacteria were analyzed by immunoblot for FLAG protein. **(C)** ClearColi and ClearColi*::dacA+* were treated (“+”) with 0.01 mM IPTG to induce expression of DacA-3xFLAG. Protein lysates from bacteria were analyzed by immunoblot for FLAG protein. **(D)** *E. coli::dacA+* were untreated or treated with increasing concentrations of IPTG to induce expression of DacA-3xFLAG. Protein lysates from bacteria were analyzed by immunoblot for FLAG protein. **(E)** *Myd88*^*-*/-^*Trif*^*-*/-^*Cgas*^*-*/-^ iBMDMs were exposed to bacteria from **Suppl. Fig. 3D** at an MOI of 50. 24 h conditioned supernatants were analyzed by ELISA for IP-10. **(F)** *Myd88*^*-*/-^*Trif*^*-*/-^*Cgas*^*-*/-^ iBMDMs were exposed to IPTG-treated *E. coli::dacA+* at increasing MOIs. 24 h conditioned supernatants were analyzed by ELISA for IP-10. **(G)** WT (circles), *Sting*^*-*/-^ (squares), *Myd88*^*-*/-^*Trif*^*-*/-^ (triangles), and *Myd88*^*-*/-^*Trif*^*-*/-^*Sting*^*-*/-^ (upside-down triangles) iBMDMs were exposed to bacteria and analyzed as in **Suppl. Fig. 2L**.

**Supplemental Figure 4.**
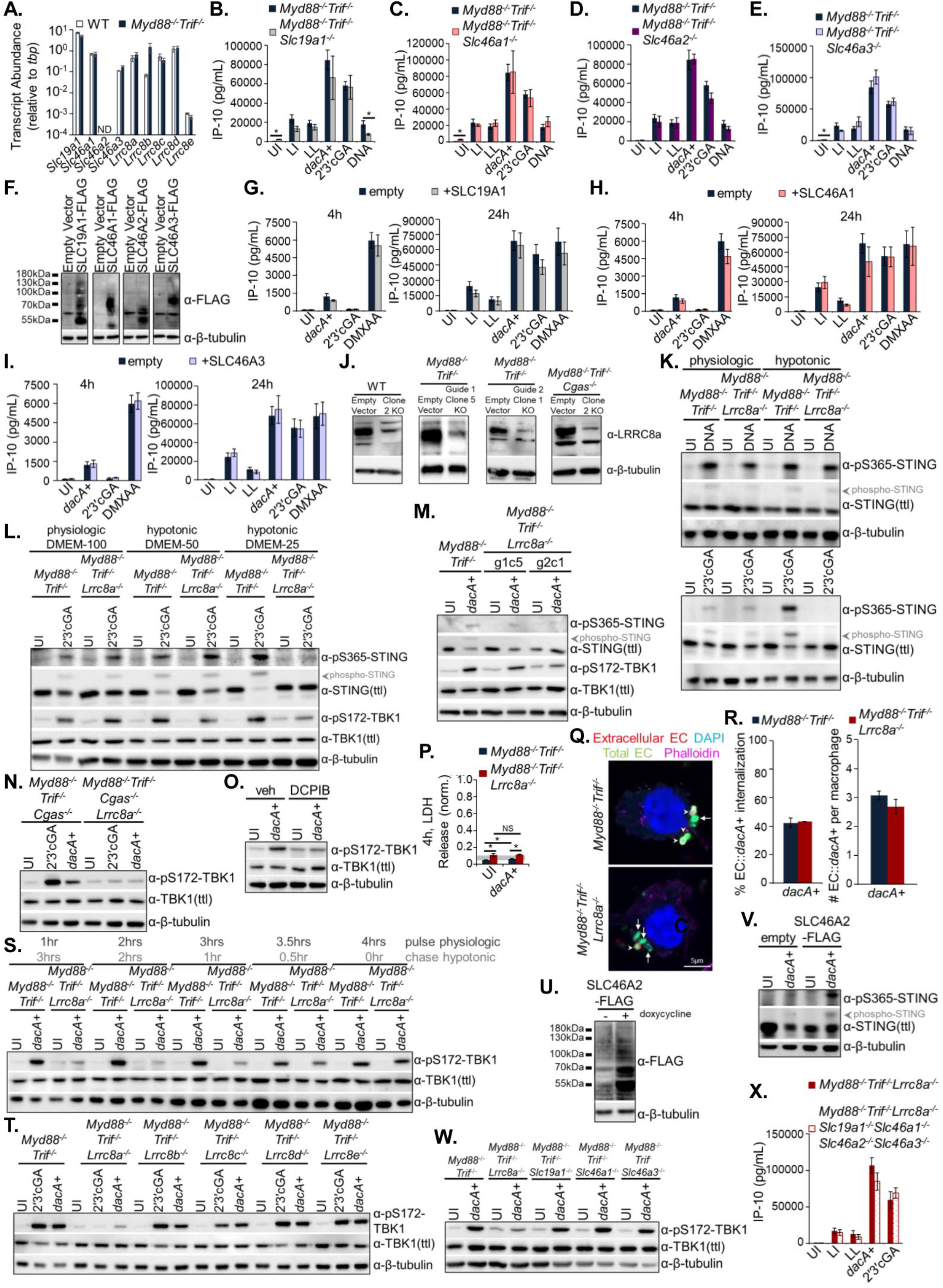
The role of CDN transporters in IFN responses to bacteria that directly stimulate STING, related to Figure 4. **(A)** RNA lysates prepared from WT and *Myd88*^*-*/-^*Trif*^*-*/-^ iBMDMs were analyzed for *Slc19a1, Slc46a1, Slc46a2, Slc46a3, Lrrc8a, Lrrc8b, Lrrc8c, Lrrc8d*, and *Lrrc8e* transcript abundance by qRT-PCR and data presented with respect to the *tbp* internal control. **(B)** *Myd88*^*-*/-^*Trif*^*-*/-^ and *Myd88*^*-*/-^*Trif*^*-*/-^*Slc19a1*^*-*/-^ iBMDMs were exposed to bacteria, including IPTG (0.01 mM)-treated *E. coli::dacA+*, at an MOI of 50, as well as 2′3′cGA (10 μg/mL), and cytosolically-delivered DNA (1 μg/mL). 24 h conditioned supernatants were analyzed by ELISA for IP-10. **(C)** *Myd88*^*-*/-^*Trif*^*-*/-^ and *Myd88*^*-*/-^*Trif*^*-*/-^*Slc46a1*^*-*/-^ iBMDMs were treated as in **Suppl. Fig. 4B**. 24 h conditioned supernatants were analyzed by ELISA for IP-10. **(D)** *Myd88*^*-*/-^*Trif*^*-*/-^ and *Myd88*^*-*/-^*Trif*^*-*/-^*Slc46a2*^*-*/-^ iBMDMs were treated as in **Suppl. Fig. 4B**. 24 h conditioned supernatants were analyzed by ELISA for IP-10. **(E)** *Myd88*^*-*/-^*Trif*^*-*/-^ and *Myd88*^*-*/-^*Trif*^*-*/-^*Slc46a3*^*-*/-^ iBMDMs were treated as in **Suppl. Fig. 4B**. 24 h conditioned supernatants were analyzed by ELISA for IP-10. **(F)** *Myd88*^*-*/-^*Trif*^*-*/-^ iBMDMs were stably reconstituted with empty vector or SLC19A1-3xFLAG, SLC46A1-3xFLAG, SLC46A2-3xFLAG, or SLC46A3-3xFLAG. Reconstituted cells were sorted for high IRES-GFP expression. Protein abundance was assessed by immunoblot for FLAG and β-tubulin. **(G)** *Myd88*^*-*/-^*Trif*^*-*/-^ iBMDMs reconstituted with empty vector or SLC19A1-3xFLAG were treated as in **Suppl. Fig. 4B**, as well as treatment with DMXAA (50 μg/mL). 4 h (left) and 24 h (right) conditioned supernatants were analyzed by ELISA for IP-10. **(H)** *Myd88*^*-*/-^*Trif*^*-*/-^ iBMDMs reconstituted with empty vector or SLC46A1-3xFLAG were treated as in **Suppl. Fig. 4G**. 4 h (left) and 24 h (right) conditioned supernatants were analyzed by ELISA for IP-10. **(I)** *Myd88*^*-*/-^*Trif*^*-*/-^ iBMDMs reconstituted with empty vector or SLC46A3-3xFLAG were treated as in **Suppl. Fig. 4G**. 4 h (left) and 24 h (right) conditioned supernatants were analyzed by ELISA for IP-10. **(J)** Protein lysates were collected from WT and *Lrrc8a*^*-*/-^ iBMDMs, *Myd88*^*-*/-^*Trif*^*-*/-^ and *Myd88*^*-*/-^*Trif*^*-*/-^*Lrrc8a*^*-*/-^ (two clones from two separate guides targeting *Lrrc8a*), and *Myd88*^*-*/-^*Trif*^*-*/-^*Cgas*^*-*/-^ and *Myd88*^*-*/-^*Trif*^*-*/-^*Cgas*^*-*/-^*Lrrc8a*^*-*/-^. Protein abundance was analyzed by immunoblot for LRRC8A and β-Tubulin. **(K)** *Myd88*^*-*/-^*Trif*^*-*/-^ and *Myd88*^*-*/-^*Trif*^*-*/-^*Lrrc8a*^*-*/-^ iBMDMs were exposed to cytosolically-delivered DNA (1 μg/mL, left blot) or 2′3′cGA (10 μg/mL, right blot) under defined hypotonic conditions (see methods). 4 h protein lysates were analyzed by immunoblot for pSTING, total STING, and β-tubulin. **(L)** *Myd88*^*-*/-^*Trif*^*-*/-^ and *Myd88*^*-*/-^*Trif*^*-*/-^*Lrrc8a*^*-*/-^ iBMDMs were exposed to 2′3′cGA (10 μg/mL) under physiologic (100% DMEM, “DMEM-100”) or hypotonic conditions (DMEM diluted with water 1:1 (“DMEM-50”) or 1:4 (“DMEM-25”)). 4 h protein lysates were analyzed by immunoblot for pSTING, total STING, pTBK1, total TBK1, and β-tubulin. **(M)** *Myd88*^*-*/-^*Trif*^*-*/-^ iBMDMs and *Myd88*^*-*/-^*Trif*^*-*/-^*Lrrc8a*^*-*/-^ iBMDM clones from **Suppl. Fig. 4J** were exposed to IPTG (0.01 mM)-treated *E. coli::dacA+* at an MOI of 50 under hypotonic conditions. 4 h protein lysates were analyzed by immunoblot for pSTING, total STING, pTBK1, total TBK1, and β-tubulin. **(N)** *Myd88*^*-*/-^*Trif*^*-*/-^*Cgas*^*-*/-^ and *Myd88*^*-*/-^*Trif*^*-*/-^*Cgas*^*-*/-^*Lrrc8a*^*-*/-^ iBMDMs were treated as in **Suppl. Fig. 4M**, with additional stimulus of 2′3′cGA (10 μg/mL). 4 h protein lysates were analyzed by immunoblot for pTBK1, total TBK1, and β-tubulin. **(O)** *Myd88*^*-*/-^*Trif*^*-*/-^ iBMDMs were treated in the absence (“vehicle”) or presence of DCPIB (20 μM) under hypotonic conditions prior to and throughout exposure to IPTG (0.01 mM)-treated *E. coli::dacA+* at an MOI of 50. 4 h protein lysates were analyzed by immunoblot for pTBK1, total TBK1, and β-tubulin. **(P)** *Myd88*^*-*/-^*Trif*^*-*/-^ and *Myd88*^*-*/-^*Trif*^*-*/-^*Lrrc8a*^*-*/-^ iBMDMs were treated as in **Suppl. Fig. 4M**. 4 h conditioned supernatants were removed and assayed for the presence of LDH as in **Suppl. Fig. 1D**. **(Q)** *Myd88*^*-*/-^*Trif*^*-*/-^ and *Myd88*^*-*/-^*Trif*^*-*/-^*Lrrc8a*^*-*/-^ iBMDMs were exposed to IPTG (0.01 mM)-treated *E. coli::dacA+* at an MOI of 10 under hypotonic conditions for 10 min. Total bacteria are labeled green and extracellular bacteria red, and the cell perimeter and nucleus were stained with phalloidin (pink) and DAPI (blue), respectively. Arrows denote intracellular bacteria. Arrowheads denote extracellular bacteria. **(R)** Percent (%) internalization of *E. coli::dacA+* (left) was determined from **Suppl. Fig. 4Q** micrographs by dividing intracellular *E. coli* by total *E. coli*, and number of *E. coli::dacA+* per macrophage (right) from **Suppl. Fig. 4Q** micrographs. **(S)** *Myd88*^*-*/-^*Trif*^*-*/-^ and *Myd88*^*-*/-^*Trif*^*-*/-^*Lrrc8a*^*-*/-^ iBMDMs were exposed to IPTG (0.01 mM)-treated *E. coli::dacA+* at an MOI of 50 under physiologic conditions for the indicated duration. Infections were then chased with hypotonic conditions for the indicated duration. 4 h total protein lysates were analyzed by immunoblot for pTBK1, total TBK1, and β-tubulin. **(T)** *Myd88*^*-*/-^*Trif*^*-*/-^, *Myd88*^*-*/-^*Trif*^*-*/-^*Lrrc8a*^*-*/-^, *Myd88*^*-*/-^*Trif*^*-*/-^*Lrrc8b*^*-*/-^, *Myd88*^*-*/-^*Trif*^*-*/-^*Lrrc8c*^*-*/-^, *Myd88*^*-*/-^*Trif*^*-*/-^*Lrrc8d*^*-*/-^, and *Myd88*^*-*/-^*Trif*^*-*/-^*Lrrc8e*^*-*/-^ iBMDMs were treated as in **Fig. 4I**, with the additional stimulus of 2′3′cGA (10 μg/mL). 4 h protein lysates were analyzed by immunoblot for pTBK1, total TBK1, and β-tubulin. **(U)** *Lrrc8a*^*-*/-^ iBMDMs stably reconstituted with a doxycycline-inducible SLC46A2-3xFLAG construct were untreated (“-”) or treated (“+”) with doxycycline (0.5 μg/mL) for 24 h and protein lysates were analyzed by immunoblot for FLAG and β-tubulin. **(V)** Reconstituted *Lrrc8a*^*-*/-^ iBMDMs from **Suppl. Fig. 4U** were exposed to IPTG (0.01 mM)-treated *E. coli::dacA+* at an MOI of 50. 4 h protein lysates were analyzed by immunoblot for pSTING, total STING, and β-tubulin. **(W)** *Myd88*^*-*/-^*Trif*^*-*/-^, *Myd88*^*-*/-^*Trif*^*-*/-^*Lrrc8a*^*-*/-^, *Myd88*^*-*/-^*Trif*^*-*/-^*Slc19a1*^*-*/-^, *Myd88*^*-*/-^*Trif*^*-*/-^*Slc46a1*^*-*/-^, and *Myd88*^*-*/-^*Trif*^*-*/-^ *Slc46a3*^*-*/-^ iBMDMs were treated as in **Suppl. Fig. 4N**. 4 h protein lysates were analyzed by immunoblot for pTBK1, total TBK1, and β-tubulin. **(X)** *Myd88*^*-*/-^*Trif*^*-*/-^*Lrrc8a*^*-*/-^ and *Myd88*^*-*/-^*Trif*^*-*/-^*Lrrc8a*^*-*/-^*Slc19a1*^*-*/-^*Slc46a1*^*-*/-^*Slc46a2*^*-*/-^*Slc46a3*^*-*/-^ iBMDMs were exposed to bacteria, including IPTG-treated *E. coli::dacA+*, at an MOI of 50, as well as 2′3′cGA (10 μg/mL). 24 h conditioned supernatants were analyzed by ELISA for IP-10.

**Supplemental Table 1.** Bacterial Strains used in this study.

| Species (Abbreviation) | Strain Designation | Source | Pathogen Status |
| --- | --- | --- | --- |
| *S. aureus* (SA) | SA113 | ATCC #35556; Gifted by David Underhill | Opportunistic pathogen |
| *S. epidermidis* (SE) | - | Gifted by Paula Watnick | Non-pathogen |
| *L. innocua* (LI) | SLCC 3423 | ATCC #33091 | Non-pathogen |
| *L. lactis* (LL) | NCTC 6681 | ATCC #19435 | Non-pathogen |
| *L. plantarum* (LP) | LP 39 | ATCC #14917 | Non-pathogen |
| *M. luteus* (ML) | ATCC 15307 | ATCC #4698 | Non-pathogen |
| *E. faecalis* (EF) | P-60 | ATCC #10100 | Opportunistic pathogen |
| *B. subtilis* (BS) | NRS 231 | ATCC #6633 | Non-pathogen |
| *E. coli* (EC) | MG1655 (K-12) | - | Non-pathogen |
| *N. perflava* (NP) | 28 | ATCC #14799 | Non-pathogen |
| *Y. pseudotuberculosis* (YP) | Plasmid-cured IP2666 with mCD1 plasmid | Gifted by Igor Brodsky | Pathogen (attenuated strain) |
| YP::dacA+ | IP26666 background | This study | Pathogen (attenuated strain) |
| *B. fragilis* (BF) | - | Gifted by Dennis Kasper | Non-pathogen |
| *B. thetaiotaomicron* (BT) | - | Gifted by Andy Goodman | Non-pathogen |
| *S. aureus* | USA300 | Gifted by Angelika Gründling | Opportunistic pathogen |
| ΔgdpP<br>*S. aureus* | USA300 background | Gifted by Angelika Gründling | Opportunistic pathogen (mutant) |
| ΔdacA<br>*S. aureus* | USA300 background | Gifted by Angelika Gründling | Opportunistic pathogen (mutant) |
| *S. aureus* | USA300 | Gifted by Victor Torres | Opportunistic pathogen |
| Δtoxins<br>*S. aureus* | USA300 background | Gifted by Victor Torres | Opportunistic pathogen (mutant) |
| *L. monocytogenes* (Lmo) | 10403S | Gifted by Darren Higgins | Pathogen |
| ΔhlyΔplcAΔplcB<br>*L. monocytogenes* | 10403S background (designation DP-L2319) | Gifted by Darren Higgins | Pathogen (attenuated strain) |

**Supplemental Table 1 continued.**
| Species (Abbreviation) | Strain Designation | Source | Pathogen Status |
| --- | --- | --- | --- |
| *S. capitis* | - | ATCC #146 | Non-pathogen |
| *S. caprae* | - | ATCC #55133 | Non-pathogen |
| *S. lugdunensis* | - | ATCC #4576 | Non-pathogen |
| *S. saprophyticus* | - | ATCC #15305 | Non-pathogen |
| *L. acidophilus* | - | ATCC #4356 | Non-pathogen |
| *L. delbrueckii* | - | ATCC #11842 | Non-pathogen |
| *E. casseliflavus* | - | ATCC #700327 | Non-pathogen |
| *E. saccharolyticus* | - | ATCC #43076 | Non-pathogen |
| *S. salivarius* | - | ATCC #13419 | Non-pathogen |
| *S. mutans* | - | ATCC #25175 | Non-pathogen |
| *S. gallolyticus* | - | ATCC #9809 | Non-pathogen |
| *P. damnosus* | - | ATCC #11842 | Non-pathogen |
| *C. pseudodiphtheriticum* | - | ATCC #10700 | Non-pathogen |
| *C. xerosis* | - | ATCC #373 | Non-pathogen |
| *K. aerogenes* | - | ATCC #13048 | Non-pathogen |
| *P. putida* | - | ATCC #12633 | Non-pathogen |
| *S. maltophilia* | - | ATCC #13637 | Non-pathogen |
| *V. natriegens* | - | ATCC #14048 | Non-pathogen |
| *S. carnosus* | - | ATCC #51365 | Non-pathogen |
| *L. rhamnosus* | strain LMS2-1 | Gifted by Dennis Kasper | Non-pathogen |
| *L. reuteri* | strain CF48-3A | Gifted by Dennis Kasper | Non-pathogen |
| *E. coli* | strain Nissle | Gifted by Dennis Kasper | Non-pathogen |
| *L. gasseri* | strain MV-22 | Gifted by Dennis Kasper | Non-pathogen |
| *L. murinus* | strain WG mouse | Gifted by Dennis Kasper | Non-pathogen |
| *L. johnsonii* | strain CB mouse | Gifted by Dennis Kasper | Non-pathogen |

**Supplemental Table 2.**
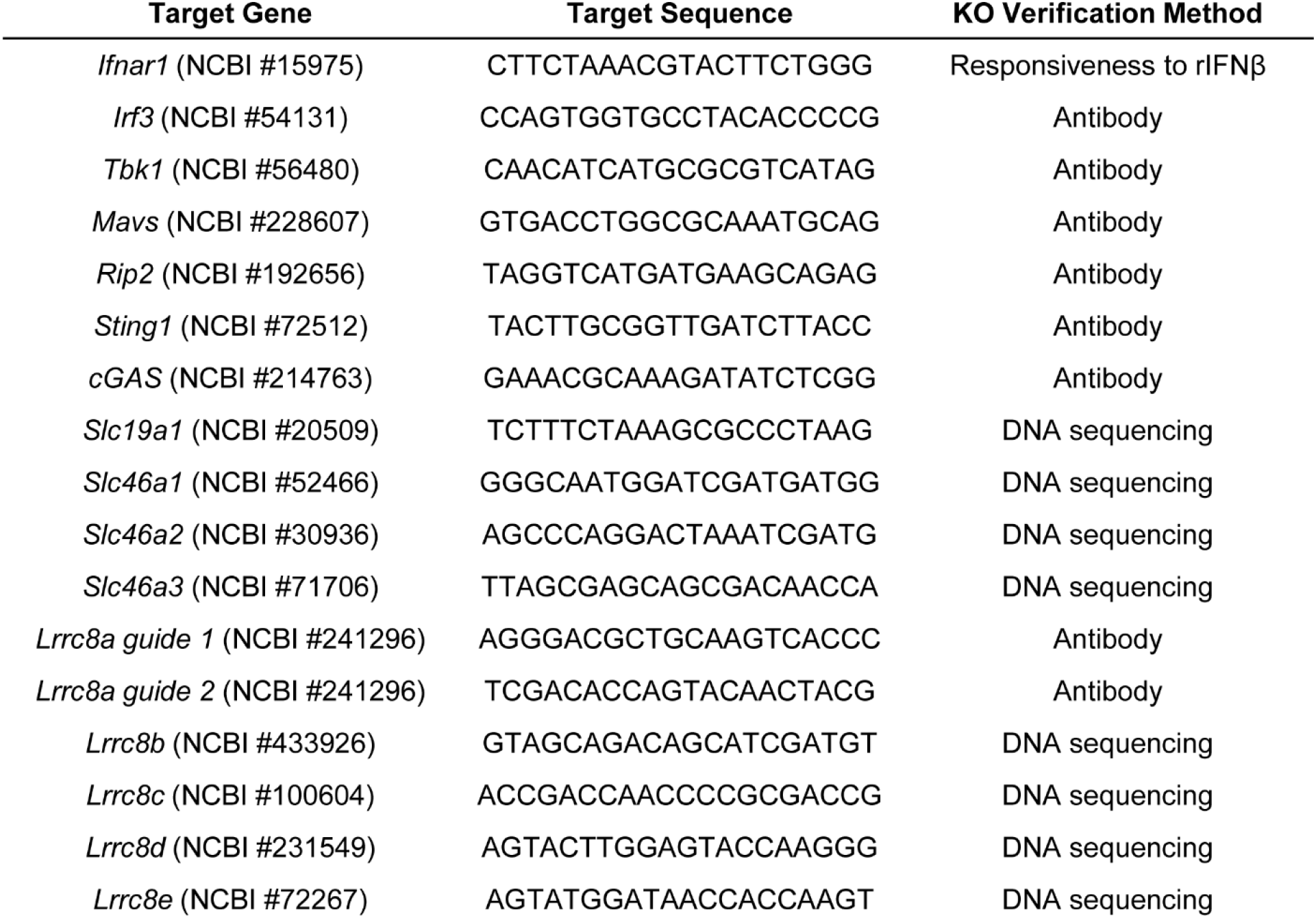
CRISPR guide target sequences for pLentiCRISPR V. 2.0 system.

